# From Sensor Design to Force Maps: A Systematic Evaluation of FRET-Based Tension Probes

**DOI:** 10.64898/2026.03.23.713753

**Authors:** Samet Aytekin, Sarah Vorsselmans, Giel Vankevelaer, Bert Poedts, Johan Hofkens, Jelle Hendrix, Susana Rocha

## Abstract

Mechanical forces transmitted through focal adhesions regulate cell behavior and disease progression, yet remain difficult to quantify at the molecular level. Genetically encoded FRET-based tension probes enable measurements of piconewton-scale forces across specific proteins in living cells, but their quantitative interpretation is highly sensitive to probe design and measurement modality. Here, we systematically compared vinculin tension sensors under identical experimental conditions, evaluating unloaded reference constructs, fluorophore pairs, mechanical sensor modules, and circularly permuted variants. Unloaded controls established a common no-force baseline and validated force-dependent readout. Among the fluorophore pairs tested, the green-red combination Clover-mScarlet-I yielded a higher unloaded FRET efficiency and hence a broader measurable dynamic range. Comparison of six mechanical sensor modules showed that the binary-response sensors FL and CC-S_2_ produced the largest force-dependent FRET changes and the steepest proximal-to-distal tension gradient. Circular permutation experiments revealed that fluorophore orientation has a strong, module-dependent influence on the measured FRET readout. Together, these results establish a comparative framework for interpreting FLIM-based vinculin tension measurements and provide practical design principles for selecting and engineering molecular tension probes.

---

Mechanical forces regulate fundamental cell behavior, including adhesion, migration, proliferation, and differentiation^1–3^ and their dysregulation contributes to pathological conditions such as fibrosis,^4^ atherosclerosis,^5^ cancer^6,7^ and others.^8^ Techniques such as traction force microscopy, micropillar arrays, and atomic force microscopy have enabled quantitative measurements of cell-generated forces at the cellular and subcellular scale,^9,10^ but they do not reveal how forces are transmitted through individual proteins.

Genetically-encoded molecular tension sensors address this limitation by measuring piconewton (pN) forces across specific proteins in living cells.^11,12^ These Förster resonance energy transfer (FRET)-based probes are composed of donor and acceptor fluorescent proteins separated by a flexible, force-sensitive linker.^13,14^ Under load, the linker extends or unfolds, increasing the donor-acceptor distance and reducing FRET efficiency. Since their development, molecular tension probes have been applied to a variety of proteins at focal adhesions, cell-cell junctions, and the nuclear envelope, enabling force measurements at distinct subcellular locations.^15^

The original vinculin tension sensor used the spring-like F40 linker to report forces in the low-pN range.^16^ Since then, sensor modules with distinct force-response characteristics have been developed, including the extension-based entropic-spring (GGSGGS)_n_,^17^ the unfolding-based HP35 domain and its stabilized variant HP35st,^18^ and the near-digital binary-response modules ferredoxin-like FL and coiled-coil CC-S_2_ (Table 1).^19,20^ These designs differ in force range, sensitivity, and opening mechanism, providing distinct strategies for probing molecular tension. Based on the range of their force-response profiles, these modules can be broadly grouped into gradual-response sensors, including F40, (GGSGGS)_n_, HP35(st), and binary-response sensors, represented by FL and CC-S_2_.

**Table 1.** Summary of common sensor modules for FRET-based molecular tension probes to measure piconewton (pN) forces over vinculin. Listed are the approximate force ranges over which each module reports tension, with the characteristic midpoint force (F_1/2_) where applicable (defined as the force at equal folded and unfolded populations for unfolding-based sensors); qualitative sensitivity based on the steepness of the force-FRET response; and the underlying opening mechanism. Contour length increase, as measured through optical tweezers experiments, denotes the increase in polypeptide contour length associated with stretching and/or unfolding, but not the end-to-end extension under physiological forces in the cell. NA, not applicable; NR, not reported; asterisk denotes estimates from published results.

|  | Force Range | F<sub>1/2</sub> | Sensitivity | Opening Mechanism | Contour Length Increase | Major Refs. |
|---|---|---|---|---|---|---|
| **F40** | 1-6 pN | NA | Low | Extension | 5-15 nm | <sup>16,17,19</sup> |
| **GGSGGS<sub>7</sub>** | 1-6 pN | NA | Intermediate* | Extension | NR | <sup>17</sup> |
| **FL** | 3-5 pN | 4 pN | High | Unfolding (Binary) | 25-30 nm | <sup>19,21</sup> |
| **HP35** | 4-9 pN* | 7.4 pN | Intermediate | Unfolding | 10 nm | <sup>18,19,21</sup> |
| **HP35st** | 8-12 pN* | 10.6 pN | Intermediate | Unfolding | 10 nm | <sup>18,19,21</sup> |
| **CC-S<sub>2</sub>** | >10 pN | 9.9 pN | High | Unfolding (Binary) | 19 nm | <sup>20</sup> |

The choice of sensor module can therefore strongly influence the measured FRET response. However, direct comparison across existing designs remains challenging because they were developed and characterized using different calibration methods, imaging conditions, FRET modalities and analysis strategies. These limitations underscore the need for a systematic, side-by-side comparison of sensor designs under identical experimental conditions.

Here, we systematically evaluated FRET-based molecular tension probes within a unified FLIM-based approach, using vinculin tension probes as a model system. We compared four key parameters: (i) force-insensitive constructs defining the unloaded (no force) state, (ii) donor-acceptor FRET pairs, (iii) sensor modules, and (iv) fluorophore orientation. Together, these experiments establish a comparative framework for interpreting vinculin tension measurements and provide practical design principles for selecting and engineering FRET-based molecular tension probes, with broader relevance to other force-bearing proteins.

## Results and Discussion

### Imaging modality shows discrepancy among force-insensitive controls

Accurate interpretation of FRET-based tension measurements requires an appropriate force-insensitive reference that defines the unloaded, high-FRET state. The widely used original vinculin tension sensor (VinTS) design served as the starting point for this analysis. It consists of the mTFP1-Venus FRET pair separated by the F40 linker, inserted between the head and tail domains (**Fig. 1a**). To define the unloaded state, we evaluated four control designs that reduce or eliminate force transmission across the sensor by distinct mechanisms. VinTL lacks the actin-binding tail domain (‘tailless’), preventing force transmission across the sensor^16,22^ (**Fig. 1b**). VinTS^I997A^ carries a point mutation that strongly reduces actin affinity, resulting in little to no tension^23–25^ (**Fig. 1c**). In VinTS-C the sensor module is fused to the C-terminus rather than inserted within vinculin, preventing mechanical loading of the module^20^ (**Fig. 1d**). Finally, the isolated tension sensor module (TSMod) diffuses freely in the cytoplasm and serves as a fully unloaded reference^16,17^ (**Fig. 1e**).

VinTS and all four force-insensitive controls were analyzed using our previously developed FLIM-phasor approach (**Supplementary Fig. 1**; **Fig. 1a-e**).^26^ Because FLIM-phasor analysis requires a donor-only lifetime (τ_D_) to define the FRET trajectory, we first examined whether mTFP1 lifetime varies with subcellular localization. Cytosolic mTFP1 (mTFP1_cyto_) exhibited a significantly longer lifetime than FA-localized mTFP1 (mTFP1_FA_; 2.71 ± 0.02 ns versus 2.56 ± 0.03 ns, respectively; **Fig. 1f**). Although the origin of this difference was not investigated here, it may reflect local variations in the molecular environment or fluorophore conformational freedom between the cytosol and FAs. Accordingly, mTFP1_cyto_ lifetime was used as τ_D_ for cytosolic TSMod, whereas mTFP1_FA_ lifetime was used for the FA-localized constructs.

**Figure 1.**
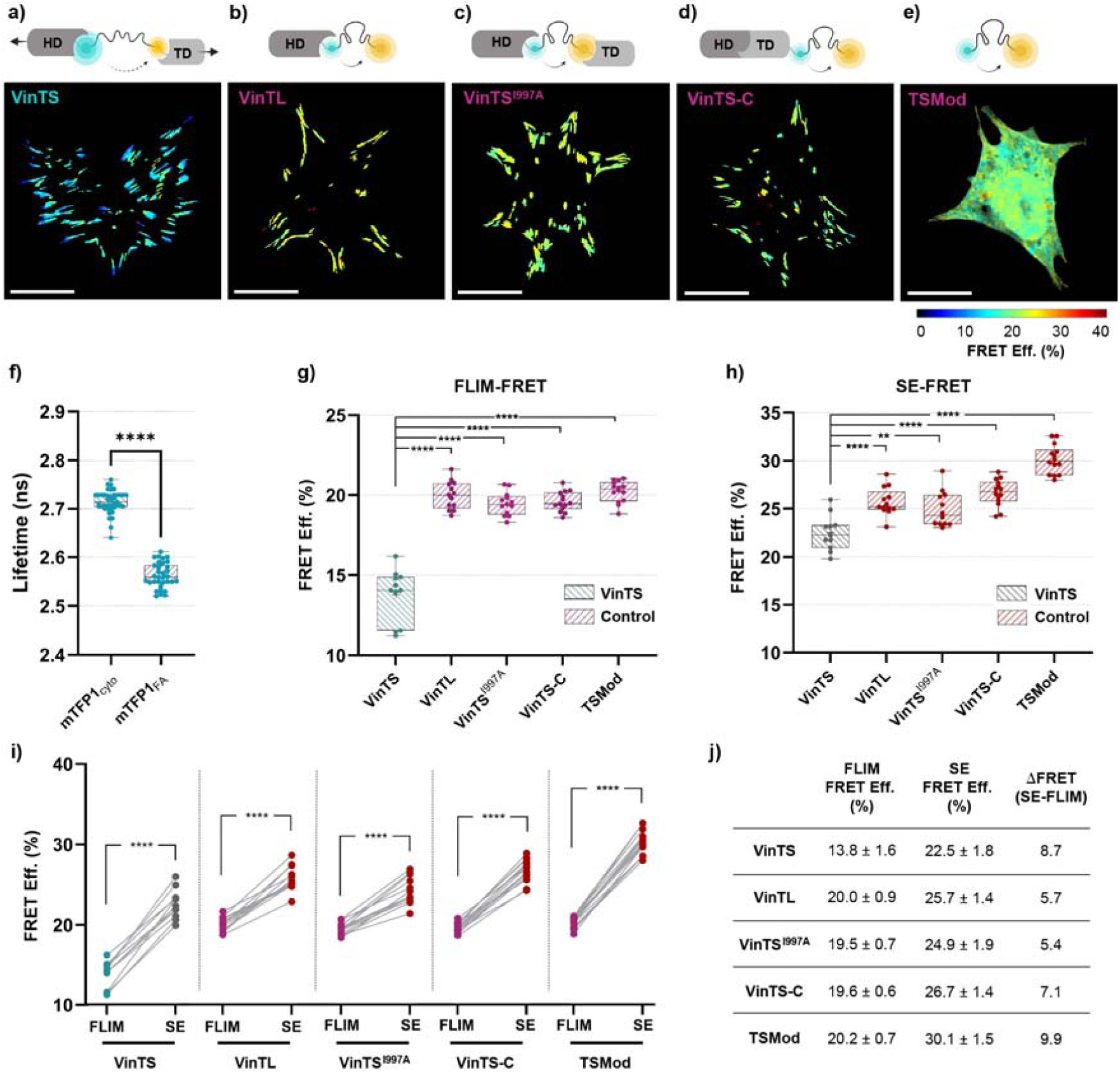
Comparison of force-insensitive controls for VinTS using FLIM and sensitized-emission FRET. (**a-e**) Schematic representations (top) and representative FRET efficiency distribution images of (**a**) tension sensor VinTS, (**b**) tailless control (VinTL), (**c**) VinTS with I997A mutation in the tail domain (VinTS^I997A^), (**d**) TSMod inserted at the C-terminus of full-length vinculin (VinTS-C), (**e**) cytosolic expression of TSMod. Scale bar: 25 µm. (**f**) Fluorescence lifetime of mTFP1 with respect to cellular localization. Lifetime values of mTFP1 localized to cytosol (mTFP1_cyto_) were compared to mTFP1 localized to FAs (mTFP1_FA_). The latter was generated by positioning mTFP1 between the vinculin head and tail domains. n = 35 and 40 cells for mTFP1_cyto_ and mTFP1_FA_, respectively. (**g**) Cell-averaged FRET comparison of VinTS (green) and force-insensitive controls (purple) using FLIM-phasor approach. (**h**) Cell-averaged FRET comparison of VinTS (gray) and force-insensitive controls (red) using sensitized-emission FRET (SE-FRET) approach. (**i**) Paired comparison of the FRET of the same cells using FLIM (green or purple) vs SE (gray or red). n = 11, 13, 12, 12 and 15 cells for VinTS, VinTL, VinTS^I997A^, VinTS-C and TSMod, respectively. Each data point represents a single cell. Each value represents the average FRET efficiency of all pixels within the FA masks for FA-localized constructs, or within the whole-cell mask for cytosolic TSMod. (j) Summary of mean FRET efficiencies measured by FLIM and SE-FRET for VinTS and the four force-insensitive controls, together with the corresponding differences (SE-FRET − FLIM). Statistical analyses between two groups were performed using unpaired two-tailed parametric Student’s t-test for (f), (g) and (h), and paired parametric Student’s t-test for (i): **p < 0.01, ***p < 0.001, ****p < 0.0001. Illustrations (a) to (e) were created using BioRender.com.

All force-insensitive controls showed significantly higher FRET efficiencies than VinTS (**Fig. 1g**), confirming mechanical loading of VinTS. Notably, all four controls showed similar FRET efficiencies by FLIM. To validate these findings with an independent FRET modality, we analyzed the same samples by intensity-based sensitized emission FRET (SE-FRET) (**Supplementary Fig. 2,** Materials and Methods). As in FLIM, all force-insensitive controls exhibited significantly higher FRET efficiencies than VinTS (**Fig. 1h**), confirming that VinTS is mechanically loaded. However, apparent FRET efficiencies were significantly higher by SE-FRET than by FLIM, consistent with discrepancies between these modalities reported by others and us.^26,27^ Importantly, in pixel-based analyses, both methods report ensemble averages over heterogeneous molecular populations and may weight these populations differently. In phasor-FLIM, low-FRET or donor-only species contribute disproportionately because of their higher photon emission, shifting apparent FRET efficiencies toward lower values.^28–30^ SE-FRET, in contrast, is derived from donor and acceptor intensities and is additionally sensitive to other factors, including donor-acceptor stoichiometry and fluorophore maturation.

Notably, the magnitude of the FLIM-to-SE shift varied across constructs (**Fig. 1i**, **Fig. 1j**). TSMod showed the largest increase, becoming the highest-FRET control under SE-FRET, whereas VinTS^I997A^ showed the smallest shift, narrowing its separation from VinTS. Thus, unlike FLIM, SE-FRET distinguished among the force-insensitive controls. One possible explanation is that the SE-FRET correction factors were derived from cytosolic calibration samples and may therefore more closely reflect the conditions experienced by cytosolic TSMod. However, this does not explain the differences among FA-localized controls, which require further investigation. These observations indicate that the choice of low- or no-tension control is modality-dependent and requires particular care for intensity-based FRET measurements.

Together, these results show that all force-insensitive constructs exhibit higher FRET than VinTS by both modalities, and can serve as a low- or no-tension reference, depending on the experimental context. For subsequent analyses, we selected VinTS^I997A^ because it best preserves the same structure and localization as VinTS.

### Fluorophore pair affects the dynamic range and robustness of FRET readout

For a given linker, higher unloaded FRET efficiency expands the measurable range available to detect force-induced changes.^17^ Fluorophore choice further affects sensor performance through differences in Förster radius, maturation kinetics, photostability, and background fluorescence. Green-red FRET pairs offer practical advantages over cyan-yellow combinations, including reduced cellular autofluorescence and lower phototoxicity.^31–33^ We therefore compared three donor-acceptor pairs in VinTS and VinTS^I997A^: the original mTFP1-Venus^16^ pair, the previously reported Clover-mRuby2 pair,^17,20,26^ and Clover-mScarlet-I, included as an alternative green-red pair given reported limitations of mRuby2 in FLIM-FRET^34–36^ (**Fig. 2a-c**). Throughout this work, constructs are referred to by their VinTS or VinTS^I997A^ backbone, with design variations indicated where relevant.

Comparison across all three FRET pairs revealed a clear performance ranking (**Fig. 2d**). Clover-mRuby2 yielded the lowest FRET efficiencies in both VinTS and VinTS^I997A^, with VinTS decreasing from 14.8 ± 2.3% for mTFP1-Venus to 11.3 ± 2.9% for Clover-mRuby2, and VinTS^I997A^ decreasing from 18.3 ± 1.0% for mTFP1-Venus to 14.8 ± 2.5% for Clover-mRuby2. This contrasts with prior SE-FRET studies in which Clover-mRuby2 outperformed the original mTFP1-Venus pair,^17^ suggesting that its reduced performance may be specific to FLIM. By contrast, Clover-mScarlet-I produced the highest FRET efficiencies, reaching 19.3 ± 1.7% for VinTS and 23.6 ± 1.3% for VinTS^I997A^. For the same sensor architecture and mechanical linker, the higher unloaded FRET observed for Clover-mScarlet-I indicates that this fluorophore pair operates closer to its Förster radius, where the FRET-distance relationship is steepest, and is therefore expected to produce a larger measurable change in FRET under force.

Because FRET variability in VinTS can reflect biological differences in molecular tension, we used VinTS^I997A^ constructs to examine fluorophore-dependent variability. In addition to its lower mean FRET, Clover-mRuby2 showed a broader FRET distribution in VinTS^I997A^ when compared with both mTFP1-Venus and Clover-mScarlet-I (**Fig. 2d**). To test whether this heterogeneity reflected imbalanced donor and acceptor expression, we quantified the donor-to-acceptor (DA) intensity ratio, defined here as the sum of donor and FRET channel intensities divided by the acceptor channel intensity (**Fig. 2e**). This DA intensity ratio depends on excitation and detection settings and was used as a relative intensity metric, rather than an absolute molar ratio. Although absolute DA ratios can differ between fluorophore pairs, they should remain relatively stable for tandem FRET sensors. Clover-mRuby2 instead exhibited a substantially broader DA ratio distribution than the other pairs. Moreover, Clover-mRuby2 showed a strong negative relationship between DA ratio and FRET efficiency, whereas mTFP1-Venus and Clover-mScarlet-I showed only weak dependence (**Fig. 2f**). Consequently, the broad DA-ratio distribution likely contributes to the broadened FRET distribution, complicating interpretation of apparent cell-to-cell variability.

To determine whether chemical fixation contributed to the lower and more spread FRET values observed with Clover-mRuby2, we compared the same VinTS^I997A^ cells before and after paraformaldehyde (PFA) fixation. Because fixation altered the donor lifetime, condition-specific donor lifetimes were used to calculate FRET efficiencies (**Supplementary Fig. 3a**). PFA fixation reduced FRET efficiency for both Clover-mRuby2 and Clover-mScarlet-I (**Supplementary Fig. 3b**), while the FRET efficiency of Clover-mRuby2 remained lower and more heterogeneous. The decrease correlated with the initial live-cell FRET efficiency for Clover-mScarlet-I, but not for Clover-mRuby2 (**Supplementary Fig. 3c**).

These results are consistent with reports that slow or incomplete chromophore maturation and pronounced photochromic behavior of mRuby2 can produce reduced and heterogenous FLIM-FRET.^26,34,35^ Non-emissive or weakly FRETing acceptors can increase the contribution of low-FRET species in photon-weighted phasor-FLIM. Overall, Clover-mScarlet-I combined the practical advantages of a green-red FRET pair with the highest unloaded FRET efficiency and robust cell-to-cell performance and was therefore selected for subsequent experiments.

**Figure 2.**
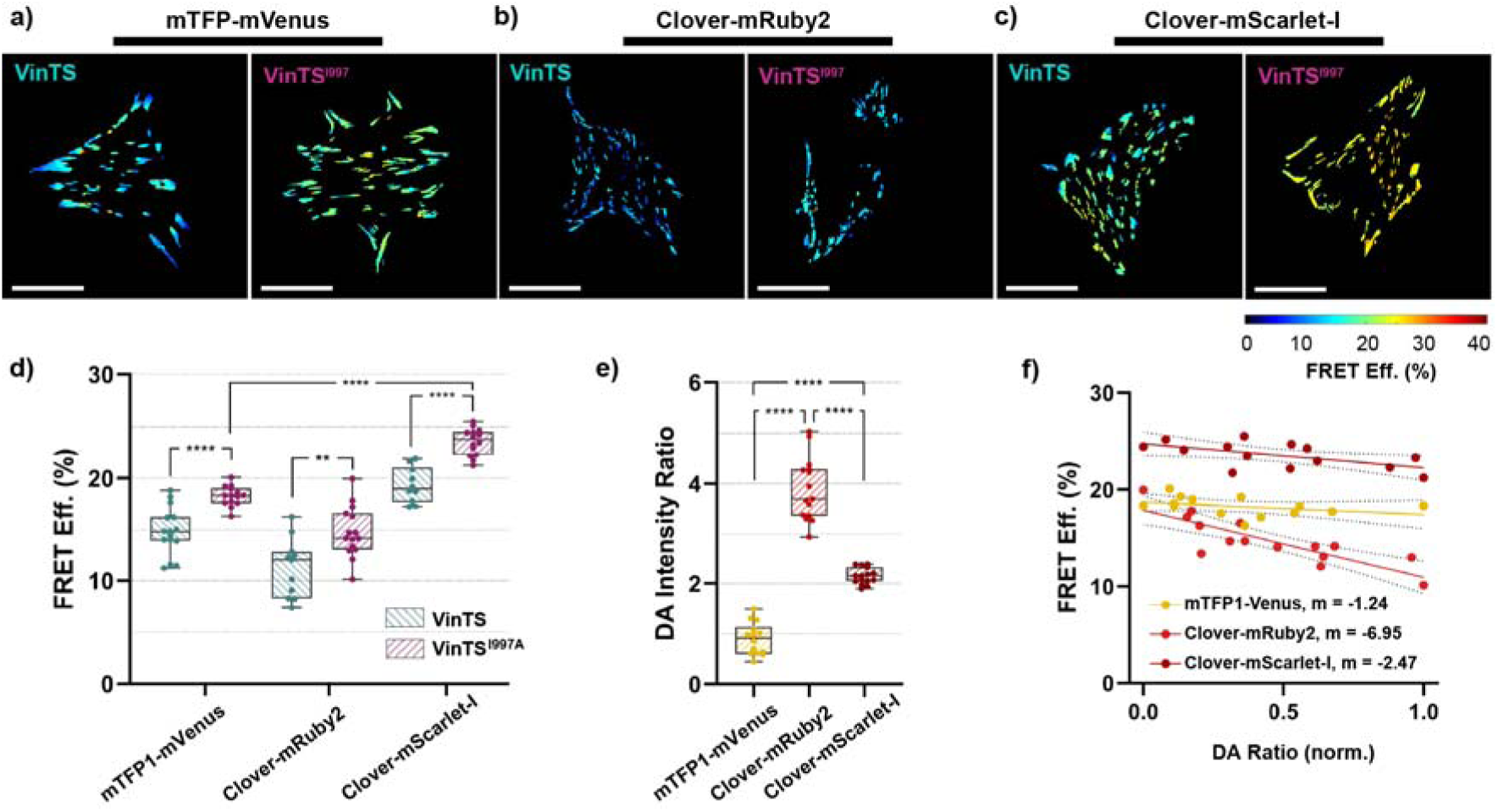
Effect of donor-acceptor fluorophore pair on the dynamic range and data heterogeneity. (**a-c**) Representative FRET efficiency distribution images of cells expressing VinTS and VinTS^I997A^ using mTFP1-Venus (a), Clover-mRuby2 (b) or Clover-mScarlet-I (c) as FRET pair. Scale bar: 25 µm. (**d**) Cell-averaged FRET efficiency comparison of VinTS (green) vs VinTS^I997A^ (purple) using mTFP1-Venus, Clover-mRuby2, and Clover-mScarlet-I FRET pairs. Each data point represents a single cell. n = 15, 11 and 10 cells for VinTS, and n = 15, 15 and 14 cells for VinTS^I997A^. (**e**) Comparison of cell-averaged donor-to-acceptor (DA) intensity ratio for VinTS^I997A^-expressing cells shown in (d) for mTFP1-Venus (yellow), Clover-mRuby2 (light red), and Clover-mScarlet-I (dark red). (**f**) Dependence of FRET efficiency on the Donor-Acceptor (DA) intensity ratio for VinTS^I997A^ cells shown in (d) and (e). Straight lines show the linear regression fit, with dashed lines representing the confidence interval of the corresponding fit. The slopes (m) of the regression lines are indicated in the figure. Statistical analyses between two groups were performed using unpaired two-tailed parametric Student’s t-test: **p < 0.01, ****p < 0.0001.

### Sensor module mechanics alter force-induced FRET response and distribution

Beyond fluorophore choice, tension-sensor performance is strongly determined by the mechanical properties of its sensor module. We therefore incorporated six mechanical linkers with distinct force sensitivities and extension or unfolding behaviors into VinTS: F40, GGSGGS□, FL, HP35, HP35st, and CC-S_2_ (**Table 1**). FRET efficiencies were compared between VinTS and the corresponding VinTS^I997A^ controls to assess module behavior under loaded and low-tension conditions (**Fig. 3a, b**).

Under low-tension conditions, HP35, HP35st, FL, and CC-S_2_ showed unloaded FRET efficiencies of ∼30%, whereas F40 and GGSGGS□ showed lower values (∼24%, **Fig. 3c, d**). This may reflect differences in donor-acceptor distance or orientation in the unloaded state. Despite these differences, all VinTS variants differed significantly from their corresponding VinTS^I997A^ controls (**Fig. 3c, d**), indicating loading within the operating range of all modules tested, including those responsive up to 10 pN.

The difference in FRET between the loaded and unloaded states was strongly module dependent, with the largest differences for the binary-response modules FL and CC-S_2_, and the smallest for F40 (**Fig. 3c, d**). This trend cannot be explained by force threshold alone: F40 and FL operate over similar low-force ranges yet show markedly different responses, whereas the higher-force CC-S_2_ exhibits the second largest FRET difference after FL. Module response therefore reflects a combination of unloaded FRET efficiency, force response range, and response mechanism (gradual vs. binary).

Because the sensor-module comparison was performed in fixed cells, we tested whether the large VinTS-VinTS^I997A^ separation for FL and CC-S_2_ was preserved in live cells. PFA fixation reduced FRET efficiency in both VinTS and VinTS^I997A^, but their separation remained significant and of similar magnitude in live and fixed conditions for both modules (**Supplementary Fig. 4a, b**). Thus, fixation alters the absolute FRET readout but does not account for the large VinTS-VinTS^I997A^ differences of the binary modules. These cell-averaged measurements establish module-dependent FRET responses but do not reveal how effectively each module resolves heterogeneous vinculin tension states.

**Figure 3.**
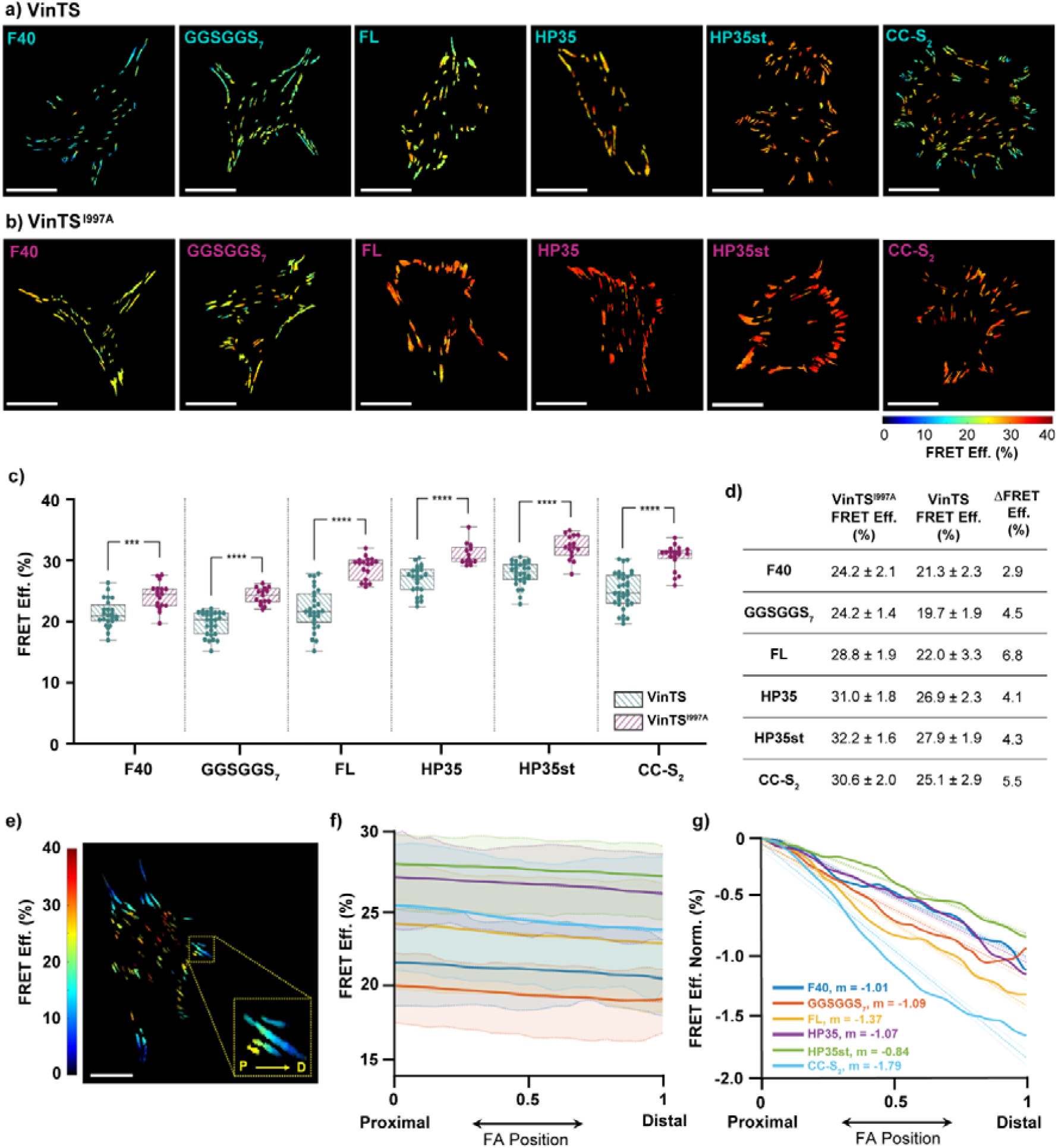
Comparison of sensor modules and sub-FA FRET gradients. (**a, b**) Representative FRET efficiency images of cells expressing VinTS (a) or VinTS^I997A^ (b) containing F40, GGSGGS_7_, FL, HP35, HP35st and CC-S_2_ as the mechanical sensing domain and Clover-mScarlet-I as fluorophore pair. Scale bar: 25 µm. (**c**) Cell-averaged FRET comparison for VinTS (green) vs VinTS^I997A^ (purple). Each data point represents a single cell. n = 24, 28, 28, 24, 29 and 35 cells for VinTS; n = 18, 15, 18, 13, 16 and 20 cells for VinTS^I997A^, respectively. Statistical analyses between two groups were performed using unpaired two-tailed parametric Student’s t-test for (c): ***p < 0.001, ****p < 0.0001. (**d**) Summary of average unloaded and loaded FRET efficiencies for each sensor module, along with their corresponding differences. (**e**) Representative cell showing the proximal-to-distal orientation and spatial FRET gradient of individual FAs. (**f**) Shaded regions represent the distribution of cell-averaged profiles and thick lines represent the global average across all cells for VinTS containing F40 (blue), GGSGGS_7_ (orange), FL (yellow), HP35 (purple), HP35st (green) and CC-S_2_ (cyan) as the mechanical sensor domain. n = 533, 796, 640, 613, 783, and 967 FAs from the same VinTS cells analyzed in (c). (**g**) Baseline-normalized profiles from (f); values indicate slopes of linear regression fits.

To examine module-dependent differences in pixel-level FRET distributions, we first compared normalized FRET efficiency distributions obtained from FA pixels (**Supplementary Fig. 5a**). Population-averaged distributions showed greater broadening of VinTS relative to VinTS^I997A^ for FL and CC-S_2_ than for the gradual-response modules. However, these distributions combine both within-cell and between-cell variability and do not directly report tension-resolving capacity. Cell-level FWHM analysis showed a significant loading-dependent increase only for GGSGGS□, with no significant differences for the other modules (**Supplementary Fig. 5b**).

Because photon statistics can also influence FRET distributions, we examined the relationship between cell-level FWHM and fluorescence intensity (**Supplementary Fig. 6a,b**). Most constructs showed no significant relation, although FWHM increased with decreasing intensity for CC-S_2_ in both VinTS and VinTS^I997A^ and for F40 in VinTS^I997A^. Thus, FRET-distribution width reflects signal- and module-dependent factors in addition to tension heterogeneity and therefore cannot, by itself, quantify tension-resolving capacity.

We therefore assessed module-dependent spatial responses using proximal-to-distal FRET profiles along individual peripheral FAs (**Fig. 3e**,**f**; individual cell profiles in **Supplementary Fig. 7a**). For comparison, profiles were baseline-normalized to a common proximal FRET value (**Fig. 3g**). Consistent with previous reports,^17,26,37^ all load-sensitive constructs showed decreasing FRET toward the distal end, whereas the corresponding VinTS^I997A^ controls showed little to no gradient (**Supplementary Fig. 7b**).

The magnitude of this response varied between modules: CC-S_2_ showed the steepest decline, followed by FL, while the gradual-response sensors showed shallower gradients. Such differences indicate that sensor-module mechanics strongly influence the spatial FRET response along FAs. The strong response of FL is consistent with its sharp binary transition near 4 pN. In contrast, F40 and GGSGGS□ showed shallower gradients despite similar force ranges, while HP35 and HP35st did not reproduce the steep CC-S2 gradient despite overlapping its force range. Thus, force range alone does not determine the spatial FRET response.

Of note, the steep CC-S_2_ gradient should not be interpreted as direct evidence of progressively higher instantaneous forces. CC-S_2_ exhibits pronounced hysteresis, refolding at substantially lower forces than it unfolds (∼8.3 pN versus ∼14.8 pN).^20^ Its low-FRET can therefore represent sensors currently under high load or sensors that unfolded previously and remain open as tension decreases. Accumulation of these history-dependent open sensors could provide a common explanation for the large loaded-unloaded FRET difference, broad population-level FRET distribution and steep sub-FA gradient observed with CC-S_2_. Thus, CC-S_2_ provides a strong readout of differences in vinculin loading, but is not a direct measure of instantaneous force magnitude.

To test whether this hysteresis produces prolonged low-FRET persistence after acute tension reduction, we monitored FL or CC-S_2_ VinTS during ROCK inhibition in live cells (**Supplementary Fig. 8a,b**). Both modules showed a rapid increase in FRET efficiency within 10 min, with comparable magnitude and kinetics. Thus, at the cell-averaged level, CC-S_2_ showed no obvious prolonged low-FRET persistence relative to FL under these conditions.

### Fluorophore orientation alters FRET readout in a module-dependent manner

Studies using FRET-based molecular tension sensors generally assume that force-induced changes in FRET primarily reflect donor and acceptor separation, with orientation effects minimized by fluorophore rotational freedom. However, linker extension or unfolding may also alter donor-acceptor geometry and thereby influence the FRET readout.

To evaluate orientation effects, we compared each module carrying the original Clover donor (Clv^0^) with its circularly permuted variant (Clv^175^), in both VinTS and VinTS^I997A^ (**Fig. 4a-b**). Circular permutation repositions the Clover termini, altering its orientation relative to the acceptor. FA-localized donor lifetime was unchanged by circular permutation (**Supplementary Fig. 9a**), supporting the interpretation that FRET differences between Clv^0^ and Clv^175^ arise from altered donor-acceptor geometry rather than donor photophysics.

Despite their lower FRET efficiency, all Clv^175^ variants retained both significant separation in FRET efficiency between VinTS and VinTS^I997A^ and proximal-to-distal FA gradients, indicating that circular permutation did not abolish tension reporting (**Supplementary Fig. 9b-e; Supplementary Fig. 10**). Gradients were shallower than for Clv^0^, but the module-dependent pattern was largely preserved.

To quantify orientation effects, we compared Clv^0^ and Clv^175^ for each sensor module using both the absolute FRET difference and the relative decrease normalized to the corresponding Clv^0^ value (**Fig. 4c, d**; **Table 2**), accounting for the differences in starting FRET efficiency between modules. In VinTS^I997A^ circular permutation reduced FRET efficiency for all modules, with absolute Clv^0^-Clv^175^ differences of 4 to 6% and relative decreases of 14.1 to 23.1% (**Table 2**). Such module dependence indicates that fluorophore orientation contributes measurably to the FRET readout and is not fully averaged out by unrestricted rotation.

For all modules, circular permutation had a smaller effect on FRET in VinTS than in the corresponding VinTS^I997A^ controls (**Fig. 4d**, **Table 2**). This is consistent with sensor loading increasing donor-acceptor separation and thereby reducing the contribution of fluorophore orientation to FRET. However, the extent of this reduction varied between modules, indicating that extension or unfolding alters donor-acceptor geometry in a strongly module-dependent manner.

**Figure 4.**
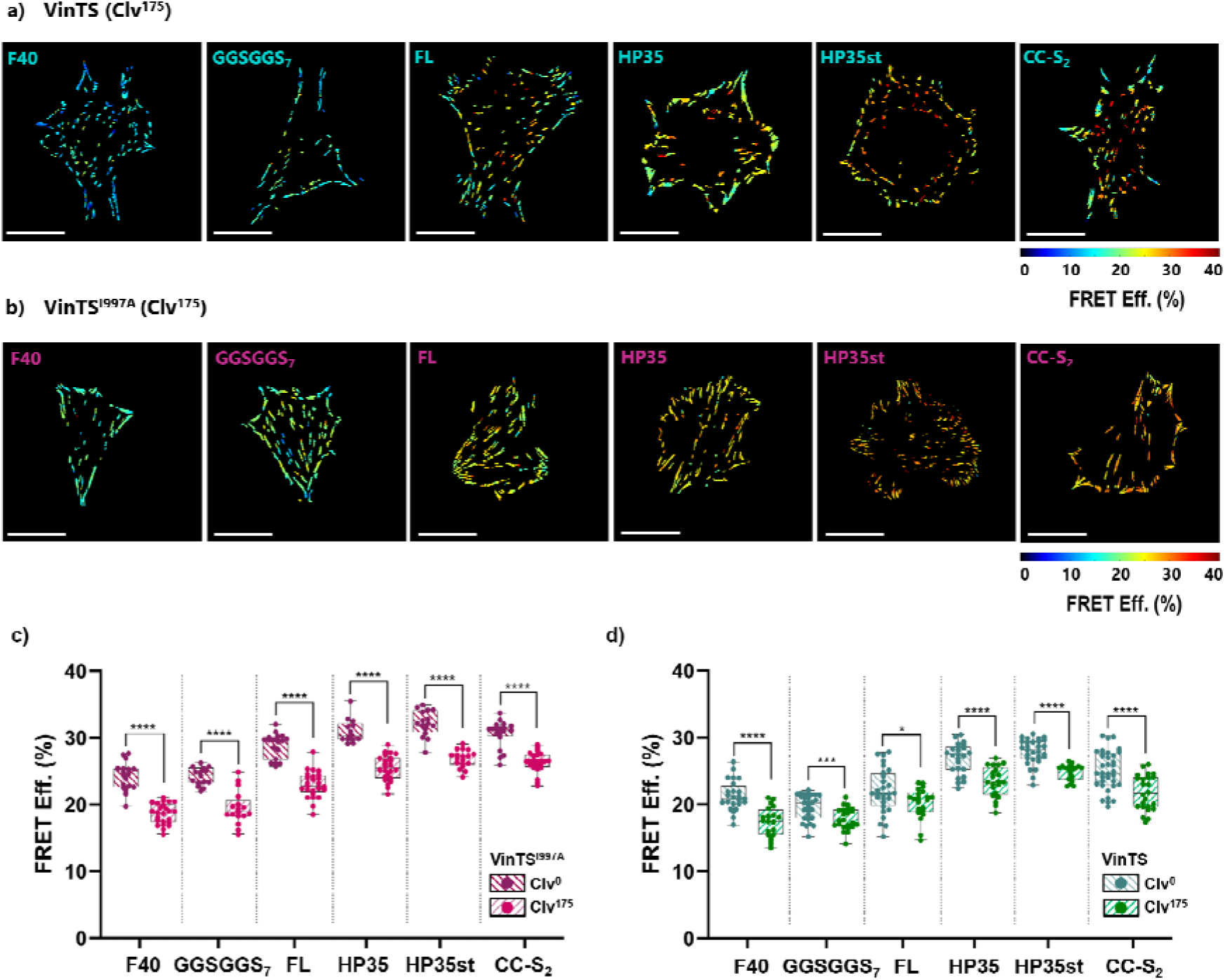
Effect of the Clover circular permutation on FRET readout of VinTS and VinTS^I997A^ across different sensor modules. (**a, b**) Representative FRET efficiency images of cells expressing VinTS (a) and VinTS^I997A^ (b) using circularly permuted Clover at residue 175 (Clv^175^) and F40, GGSGGS_7_, FL, HP35, HP35st or CC-S_2_ as the mechanical sensor domain. Scale bar: 25 µm. (**c, d**) Cell-averaged FRET efficiency comparison of non-permuted Clv^0^ and permuted Clv^175^ variants for VinTS^I997A^ (c) and VinTS (d). For VinTS (Clv^175^), n = 20, 23, 21, 21, 18, and 24 cells, respectively; for VinTS^I997A^ (Clv^175^), n = 24, 17, 25, 24, 18, and 24 cells, respectively. Clv datasets are from Fig. 3c. Each data point represents a single cell. Statistical analyses between two groups were performed using unpaired two-tailed parametric Student’s t-test: *p < 0.05, ***p < 0.001, ****p < 0.0001.

**Table 2.** Absolute and relative FRET efficiency changes upon Clover circular permutation. Values show average FRET efficiencies of Clv^175^ VinTS^I997A^ and VinTS constructs. FRET eff. (%) decrease represents the absolute difference between Clv and Clv¹ FRET efficiencies. Relative FRET change is defined as the FRET efficiency decrease normalized to the corresponding Clv value, expressed as a percentage.

|  | **FRET eff. (%) VinTS<sup>I997A</sup>** | **FRET eff. (%) VinTS** | **FRET eff. (%) decrease VinTS<sup>I997A</sup>** | **Relative FRET change (%) VinTS<sup>I997A</sup>** | **FRET eff. (%) decrease VinTS** | **Relative FRET change (%) VinTS** |
|---|---|---|---|---|---|---|
| **F40** | 18.6 ± 1.5 | 17.4 ± 2.2 | 5.6 | 23.1 | 4.0 | 18.5 |
| **GGSGGS<sub>7</sub>** | 19.5 ± 2.5 | 17.8 ± 1.7 | 4.7 | 19.4 | 1.8 | 9.4 |
| **FL** | 23.0 ± 2.0 | 20.2 ± 2.3 | 5.8 | 20.3 | 1.8 | 8.3 |
| **HP35** | 25.5 ± 1.9 | 23.4 ± 2.3 | 5.5 | 17.8 | 3.5 | 12.9 |
| **HP35st** | 26.8 ± 1.3 | 24.8 ± 1.2 | 5.3 | 16.6 | 3.1 | 11.0 |
| **CC-S<sub>2</sub>** | 26.3 ± 1.5 | 21.8 ± 2.6 | 4.3 | 14.1 | 3.3 | 13.2 |

CC-S□ showed similar relative effects of circular permutation in both VinTS^I997A^ and VinTS (14.1% and 13.2%, respectively; **Table 2**). Given its ∼19 nm contour-length increase upon unfolding, the unfolded population may contribute little to FRET, leaving the measured response dominated by closed sensors (**Fig. 5**, left). Alternatively, if unfolded CC-S remains within the FRET-sensitive range, constrained donor–acceptor geometries may face a similar relative decrease upon circular permutation in both states. The remaining modules showed smaller relative changes in VinTS than in VinTS^I997A^, with the strongest attenuation observed for FL and GGSGGS□ (**Table 2**). Unlike the large-separation scenario, these sensors still contribute to FRET in their loaded configuration while showing reduced sensitivity to circular permutation. Such attenuation may reflect either an increased orientational sampling upon loading, arising either from greater freedom of individual fluorophores or from heterogeneous load-dependent orientations averaged within each population (**Fig. 5**, right).

An additional consideration is that forces insufficient to trigger unfolding may still perturb the folded module and constrain the fluorophores toward preferred, non-isotropic orientations. Such pre-unfolding reorientation could affect VinTS and VinTS^I997A^ differentially, thereby changing the relative effect of circular permutation between them. This possibility is particularly relevant for structurally constrained modules: unlike the flexible, unstructured GGSGGS□ linker, HP35, HP35st, and CC-S_2_ form helical structures that may restrict fluorophore rotational freedom and transmit load-induced conformational changes to donor-acceptor orientation.^38^

Overall, these results show that fluorophore orientation contributes substantially to the FRET readout in both unloaded and loaded states, indicating that donor and acceptor are not freely rotating in these sensor modules. Thus, interpreting force-induced FRET changes requires considering load-dependent changes in donor-acceptor distance and orientation, with their relative contributions depending on the sensor architecture.

**Figure 5.**
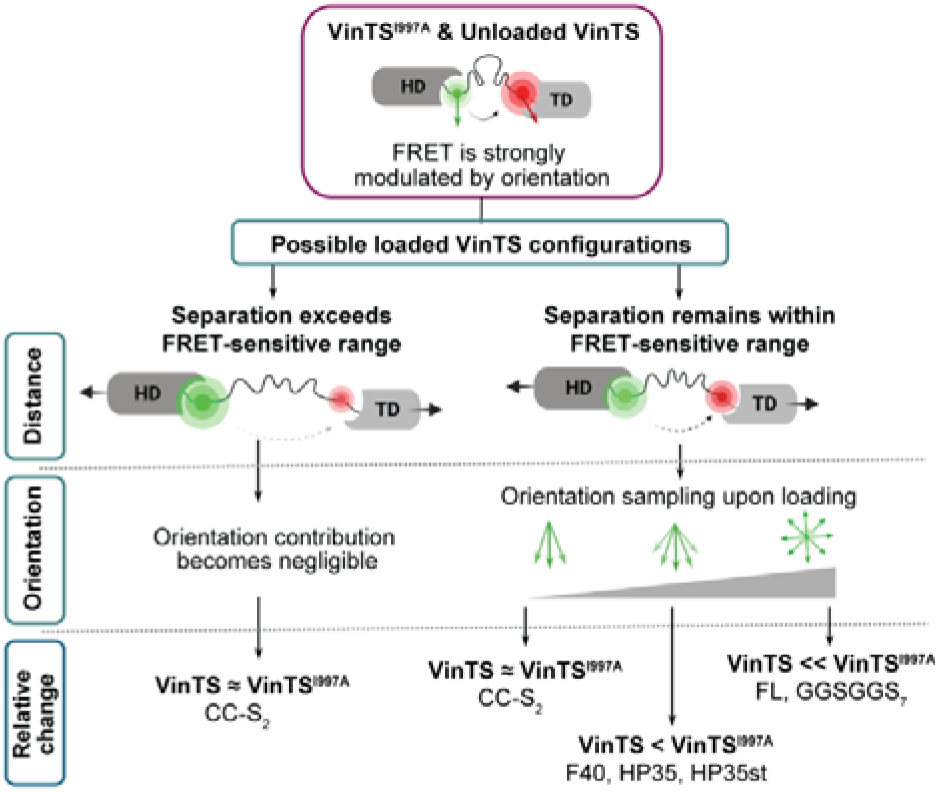
Proposed model for the module-dependent effect of fluorophore orientation on VinTS FRET readout. In the unloaded VinTS^I997A^ state, or in folded VinTS molecules, donor-acceptor separation remains small and FRET is strongly influenced by fluorophore orientation. Upon loading, sensor opening can increase donor-acceptor separation beyond the FRET-sensitive range, making orientation effects negligible. Alternatively, donor-acceptor separation remains within the FRET-sensitive range, where increased orientation sampling upon sensor opening reduces the effect of circular permutation in loaded VinTS, thereby increasing the difference between the relative FRET changes measured in VinTS and VinTS^I997A^.

## Conclusion

FRET-based molecular tension probes provide access to pN-scale forces across specific proteins, but their readout depends strongly on sensor design and measurement modality. Here, by systematically comparing key design variables within a unified FLIM-based framework, we establish how fluorophore pair, sensor-module mechanics, and fluorophore orientation shape vinculin tension measurements.

Clover-mScarlet-I provided the highest unloaded FRET and the most robust green-red FLIM-FRET readout among the fluorophore pairs tested. Across six mechanical modules, the binary-response modules FL and CC-S_2_ provided the largest differences between loaded and force-insensitive constructs, with CC-S_2_ also exhibiting the steepest proximal-to-distal sub-FA gradient. However, CC-S□ hysteresis means that its low-FRET state can reflect force history as well as current loading, preventing direct assignment of its FRET response to instantaneous force magnitude. Higher-threshold binary sensors with minimal hysteresis could help define the upper range of vinculin loading more precisely. Circular permutation further revealed module-dependent orientation effects, demonstrating that FRET changes cannot be interpreted solely in terms of donor-acceptor distance.

The pronounced responses of FL and CC-S□ do not make binary-response sensors preferable for every application. Calibrated gradual-response sensors provide a continuous force-dependent readout and are better suited to estimating force magnitude within their operating range, whereas binary-response sensors can potentially report the fraction of molecules undergoing force-induced unfolding when their folded and unfolded states can be distinguished. Sensor selection should therefore depend on the type of force information required.

Together, these findings provide practical principles for selecting and interpreting molecular tension probes. Accounting for fluorophore properties, linker mechanics, sensor geometry, and measurement modality will be important when designing and interpreting tension probes for vinculin and other force-bearing proteins.

## Materials and Methods

### Cell culture and transient expression

Vinculin-/-mouse embryonic fibroblasts (MEF-/-), kindly provided by Wolfgang H. Goldmann and Ben Fabry,^39^ were cultured in Dulbecco’s Modified Eagle Medium (DMEM) containing 4.5 g/L D-glucose (Gibco) and supplemented with 10% (v/v) fetal bovine serum (Sigma-Aldrich), 1% GlutaMax (Thermo Fisher), 0.1% gentamycin (Carl Roth), 1% (v/v) sodium pyruvate (Sigma-Aldrich), and 1% (v/v) non-essential amino acids (Sigma-Aldrich). Cells were maintained at 37 °C and 5% CO□, and passaged at 80-90% confluency. For microscopy imaging, cells were seeded at 5 × 10□ cells cm□² on 8-well plates coated overnight with 10 µg/mL fibronectin at 4 °C. After 24 h, cells were transfected using TransIT-X2 (Mirus Bio, MIR 6006) according to the manufacturer’s protocol. Briefly, 300 ng plasmid DNA and 1 µL TransIT-X2 were diluted in 100 µL serum-free DMEM per well, incubated for 15 min at room temperature, and added dropwise to the cells. Cells were allowed to express the constructs overnight and fixed the following day with 4% paraformaldehyde (PFA, Thermo Scientific). For paired live-versus-fixed measurements, cells were imaged live, fixed with 4% PFA for 30 min, and the same cells were re-imaged. For ROCK inhibition experiments, cells were maintained in HBSS at 37 °C and treated with 5 µM Y-27632 for up to 30 min.

### Molecular cloning

VinTS, VinTL, TSMod containing the mTFP1-F40-Venus module, and Vinculin-Venus were gifts from Martin Schwartz (Addgene #26019, #26020, #26021, #27300),^16^ and VinTS^I997A^ was a gift from Brenton Hoffman (Addgene #111828).^40^ VinTS-C was generated by PCR amplification of the mTFP1-F40-Venus module with 5′ EcoRI and 3′ XbaI restriction sites and replacement of Venus in the Vinculin-Venus backbone.

To enable modular exchange of fluorophores and module linkers, a VinTS master plasmid was generated by introducing unique restriction sites: XhoI between the vinculin head domain and donor fluorophore, Kpn2I between donor and linker, BamHI between linker and acceptor, NotI between acceptor and vinculin tail, and EcoRI downstream of the stop codon. A TSMod master plasmid lacking the vinculin head and tail domains was generated by ligating the module region (XhoI-EcoRI) into a pcDNA3.1 backbone PCR amplified with matching restriction sites. A VinTS^I997A^ master plasmid was generated by replacing the vinculin tail domain with the I997A mutant via NotI-EcoRI digestion and ligation.

mTFP1 was replaced by Clover or circularly permuted Clover (Clv^175^) using XhoI-Kpn2I, and Venus by mRuby2 or mScarlet-I using BamHI-Bsp1407I. To facilitate acceptor exchange, a truncated Clover variant lacking the C-terminal MDELYK residues and the corresponding Bsp1407I site was used throughout. All linker modules, including the codon-optimized F40 and GGSGGS_7_, were synthesized as double-stranded DNA fragments (gBlocks, Integrated DNA Technologies) flanked by 5′ Kpn2I and 3′ BamHI sites.

Cytosolic Clover (Clv_cyto_) was a gift from Michael Lin (Addgene #40259).^31^ Cytosolic mTFP1 (mTFP1_cyto_) and Venus were generated by replacing Clover with mTFP1 or Venus via EcoRI-Bsp1407I digestion. Clv^175^ was constructed by amplifying residues 1-175 with a 5′ GGTGGS linker and 3′ Kpn2I site and residues 176-236 with a 5′ XhoI site and 3′ GGTGGS linker, followed by overlap-extension PCR and insertion via XhoI-Kpn2I. FA-targeted donor constructs (mTFP1_FA_, Clv^0^ and Clv^175^_FA_) were generated by replacing the module region in the VinTS master plasmid with the corresponding fluorophore using XhoI-NotI.

For SE-FRET calibration, a high-FRET control was generated by PCR amplification of mTFP1 and Venus with complementary GGSGGS linkers followed by Kpn2I-mediated self-ligation. The TSMod master plasmid module was then replaced with the resulting mTFP1-(GGSGGS)_2_-Venus construct using XhoI-EcoRI. A low-FRET control (mTFP1-TRAF-Venus) was generated by replacing the sensor linker in TSMod with the TRAF domain amplified from Cerulean-TRAF-Venus, a gift from Steven Vogel (Addgene #27803),^41^ using Kpn2I-BamHI.

All constructs were verified by Sanger sequencing, with complete sequences provided in the Source Data. Primer and gBlock sequences are provided in **Supplementary Tables 1 and 2**, respectively.

### Fluorescence imaging

FLIM-FRET imaging was performed on a Leica TCS SP8 confocal microscope equipped with the FALCON (FAst Lifetime CONtrast) module. Images were collected using a 63× HC PL APO water-immersion objective (NA 1.20) at 512 × 512 pixels, corresponding to a field of view of 92.26 µm and a pixel size of 180 nm. Scanning was performed unidirectionally at 50 Hz line scan frequency with a pixel dwell time of 30 µs. mTFP1 was excited by two-photon excitation at 880 nm with a repetition rate of 80 MHz, and emission was detected between 460-500 nm using a HyD SMD detector. Venus was excited at 514 nm, with emission collected between 520-630 nm using a HyD detector. Clover and its circularly permuted variant were excited at 488 nm, and fluorescence was detected between 495-540 nm using a HyD SMD detector. mRuby2 and mScarlet-I were excited at 561 nm, with emission collected between 570-740 nm using a HyD detector. For FLIM vs. SE-FRET and FRET-pair experiments, excitation was performed at 80 MHz, whereas sensor module and fluorophore experiments were acquired at 20 MHz. For SE-FRET, mTFP1 and Venus were recorded sequentially using the same excitation and detection settings with two detectors used simultaneously to acquire donor, FRET, crosstalk, and acceptor channels.

### FLIM-FRET analysis

For FRET analysis, pixel-wise fluorescence decay curves were converted into phasor space using the integrated Phasor module of the SP8 FALCON system. Briefly, the measured decay ***I(t)*** was Fourier transformed to obtain the real (***G***) and imaginary (***S***) components:

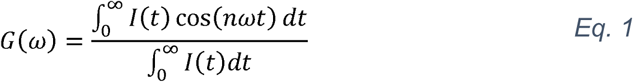

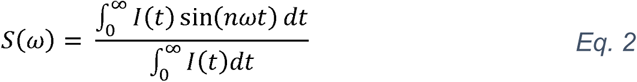

where ***n*** denotes the selected harmonic (here, 1), ***f*** is the laser repetition rate (80 or 20 MHz) and ω is 2πf. The resulting ***G*** and ***S*** coordinates generate a two-dimensional phasor histogram representing the pixel-wise lifetime distribution. Phasor coordinates were exported and processed using our previously developed custom MATLAB script for calculation of phasor-based FRET efficiencies.^26^ The donor-only lifetime position defined the origin of the FRET efficiency trajectory, while the endpoint was set to an approximate background position (G = 0.5, S = 0.25) determined from mock-transfected cells. Each pixel was assigned the FRET efficiency corresponding to the nearest position along the trajectory, yielding a spatially resolved FRET map. To reduce noise and smooth the resulting FRET efficiency distribution, a 3 × 3 pixel median filter was applied to the final FRET image.

### Sensitized-emission FRET Analysis

SE-FRET measurements of (mTFP1-F40-Venus)-based VinTS, VinTL, VinTS^I997A^, VinTS-C and TSMod were performed as described by Gates et al.^42^ Four channels were acquired: i) donor excitation with donor emission (I_DD_, donor channel), ii) donor excitation with acceptor emission (I_DA_, FRET channel), iii) acceptor excitation with acceptor emission (I_AA_, acceptor channel), and iv) acceptor excitation with donor emission (I_AD_, crosstalk channel). The crosstalk signal was consistently minimal and therefore neglected in subsequent analysis.

The corrected FRET signal (F_c_) was calculated as:

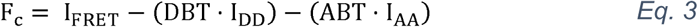

where DBT denotes donor bleedthrough, and ABT accounts for direct acceptor excitation at the donor excitation wavelength. I_DD_, I_AA_, and I_FRET_ correspond to the donor, acceptor, and FRET channel intensities, respectively.

DBT was determined from donor-only cells by binning pixels by donor intensity (I_DD_) and averaging the I_FRET_ values within each bin. DBT was defined as the slope of a linear fit of I_FRET_ versus I_DD_. ABT was calculated analogously from acceptor-only cells using I_FRET_ plotted versus I_AA_.

FRET efficiency (E) was computed from Fc as:

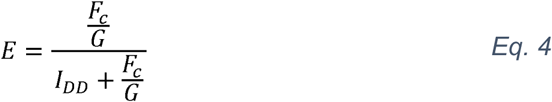

where G is a proportionality constant correcting differences in fluorophore brightness and detection efficiency between donor and FRET channels. G-factor was determined from a high-FRET fusion construct (mTFP1-GGSGGS -Venus) and a low-FRET fusion construct (mTFP1-TRAF-Venus). For each construct, acceptor-normalized corrected FRET (F_c_/I_AA_) and donor intensity (I_DD_/I_AA_) were assembled into two-dimensional histograms, and G was defined as the slope of the line connecting the modes of the two distributions:

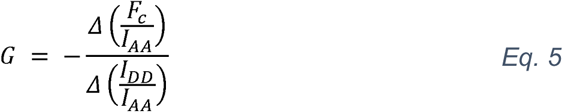

FRET efficiencies were calculated per pixel and averaged over the FA mask for FA-localized constructs or the whole-cell mask for cytosolic TSMod before comparison with FLIM-derived FRET values.

Relative donor-to-acceptor (DA) intensity ratios for different FRET pairs were calculated as:

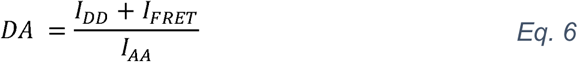

### Sub-FA FRET gradient analysis

For FA segmentation, binary masks were generated from donor images using the Ilastik machine-learning-based segmentation tool.^43^ FRET efficiency profiles along the individual FAs were analyzed using our previously developed custom MATLAB code.^26^ For each FA, the long axis was identified, and a line drawn through the FA centroid was extended to intersect the FA boundaries. The intersection point closer to the cell centroid was defined as the proximal end. FRET efficiencies were sampled along lines perpendicular to the long axis using an iterative pixel-based averaging procedure. The resulting profile was linearly interpolated to a normalized axis of 50 points. FAs shorter than 10 pixels were excluded to avoid discontinuities during interpolation.

### Statistics and reproducibility

All experiments were performed at least three independent times, with technical replicates and biologically independent samples as indicated. Experiments were not randomized, and investigators were not blinded during data acquisition or analysis. Data quantification was performed using automated analysis pipelines. Graphs were prepared using MATLAB R2024b or GraphPad Prism 10.4.1 (GraphPad Software). Source data contain the underlying data points for all graphs and the corresponding statistical analyses. Data normality was evaluated using the Shapiro-Wilk test. Normally distributed data were analyzed using two-tailed unpaired Student’s *t*-tests, whereas the Mann-Whitney *U* test was applied for non-normally distributed data. Statistical significance was defined as *p < 0.05, **p < 0.01, ***p < 0.001, ****p < 0.0001. Relationships between two variables were assessed using simple linear regression with 95% confidence intervals. Additional statistical details are provided in the figure legends.

## Data Availability

Plasmids generated in this study will be deposited in Addgene and made available upon publication. Raw microscopy data and source data files are available from the corresponding author upon request. The analysis code used for FLIM-FRET, sensitized-emission FRET (SE-FRET), population-level FRET histogram analysis, and sub-focal-adhesion FRET gradient analysis were adapted from previously published codes.^26^

## Supporting Information Available

The following files are available free of charge.

*Aytekin_et_al_ACS_Sensors_Supporting_Information.pdf.* Supplementary FLIM and SE-FRET analyses; live-versus-fixed validation of Clover-mRuby2 and Clover-mScarlet-I and of FL- and CC-S -based tension sensors; cell-level FRET-distribution and FWHM analyses; fluorescence-intensity dependence of FWHM; individual-cell sub-FA FRET profiles; Y-27632 tension-relaxation experiments; circular-permutation sub-FA analyses; primer sequences; and sensor-module sequences (PDF).

## Notes

A preprint of this article was posted on bioRxiv: Aytekin, S.; Vorsselmans, S.; Vankevelaer, G.; Poedts, B.; Hofkens, J.; Hendrix, J.; Rocha, S. From Sensor Design to Force Maps: A Systematic Evaluation of FRET-based Vinculin Tension Sensors. bioRxiv **2026** March, 2026.03.23.713753. DOI: 10.64898/2026.03.23.713753 (accessed November 10, 2026).

## Acknowledgments

The authors thank Rik Nuyts for assistance with microscopy measurements and imaging optimization, and Débora Linhares for her continued technical support with cell culture. This work was supported by KU Leuven internal funding (C14/22/085) and Research Foundation Flanders (FWO) through projects with grant numbers G0C2422N, G0A8L24N and G0B9922N. In addition, S.A. is the recipient of a FWO fellowship (1S95125N) and S.V. acknowledges KU Leuven Facultaire Luik Onderzoeksfonds (FLOF) for financial support.

---

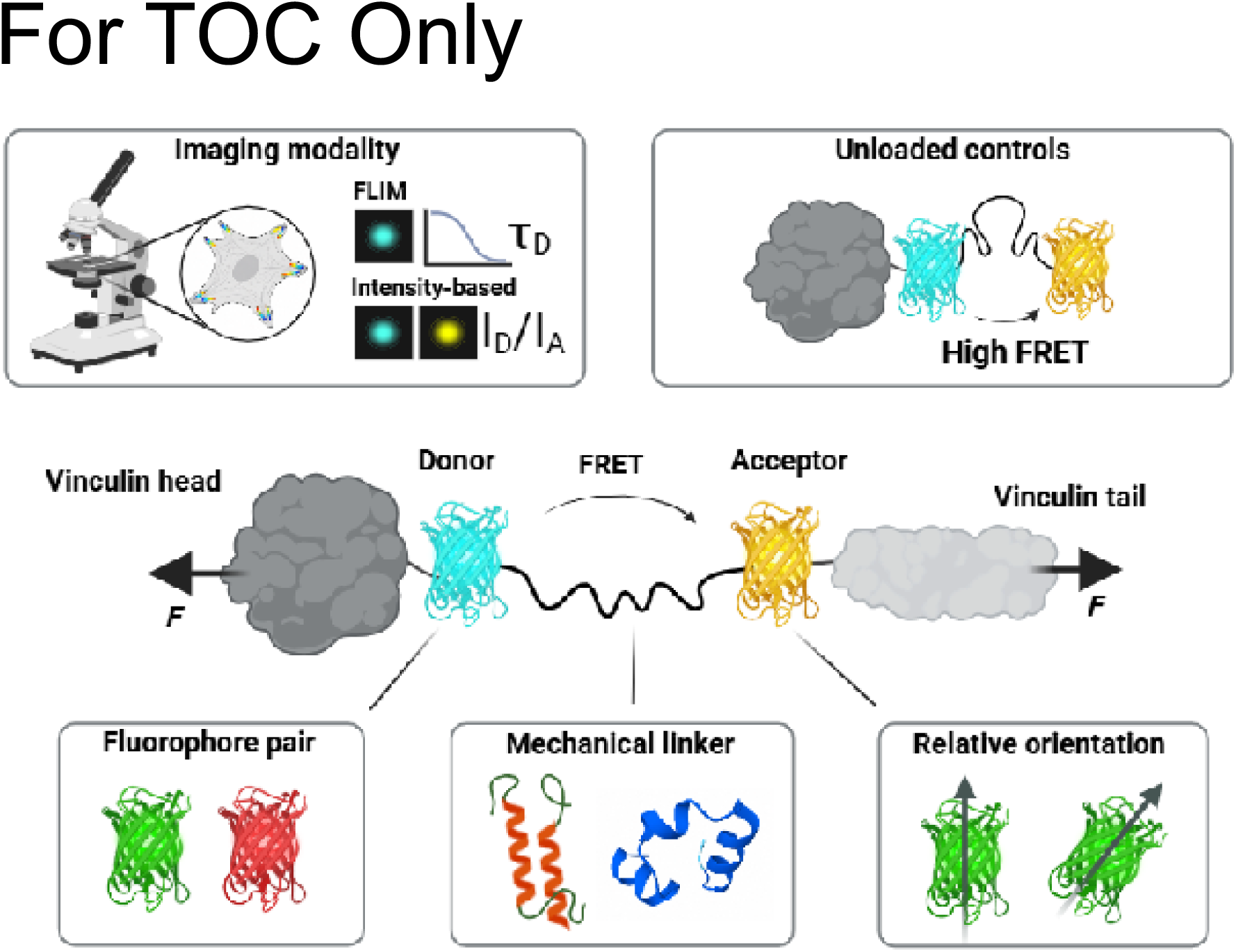

