## Supporting Information for "From Sensor Design to Force Maps: A Systematic Evaluation of FRET-Based Tension Probes"

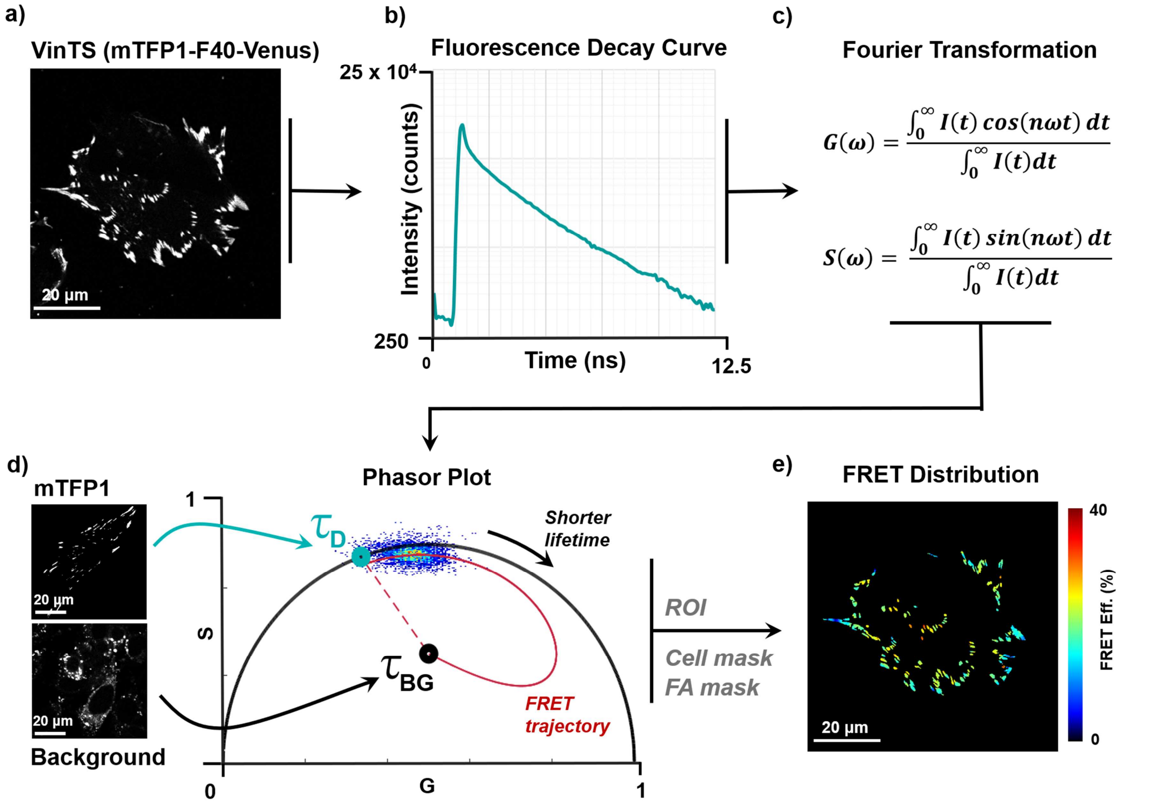


Supplementary Figure 1. FLIM-phasor approach to calculate the FRET efficiency of molecular tension probes. (a) Representative donor channel image of a cell expressing VinTS with F40 sensor module and mTFP1-Venus as FRET pair. (b) Pixel-averaged fluorescence decay curve of the donor channel acquired using Leica SP8 FALCON time-domain FLIM acquisition module with 80 MHz pulse repetition rate. (c) Fourier transformation of the decay curve into the phasor coordinates G and S, computed as the normalized first harmonic of the cosine and sine transforms. (d) Phasor plot showing the donor-only lifetime position ($\tau_D$) obtained from cells expressing FA-localized mTFP1, and the background position ($\tau_{BG}$) obtained from non-transfected cells. The line connecting $\tau_D$ and $\tau_{BG}$ defines the FRET trajectory. Phasor point of each pixel of the image is then projected to the closest point on the trajectory to obtain the FRET efficiency value. (e) Final pixel-level FRET efficiency distribution image generated after manually selecting the region of interest (ROI), applying cell and FA masks.

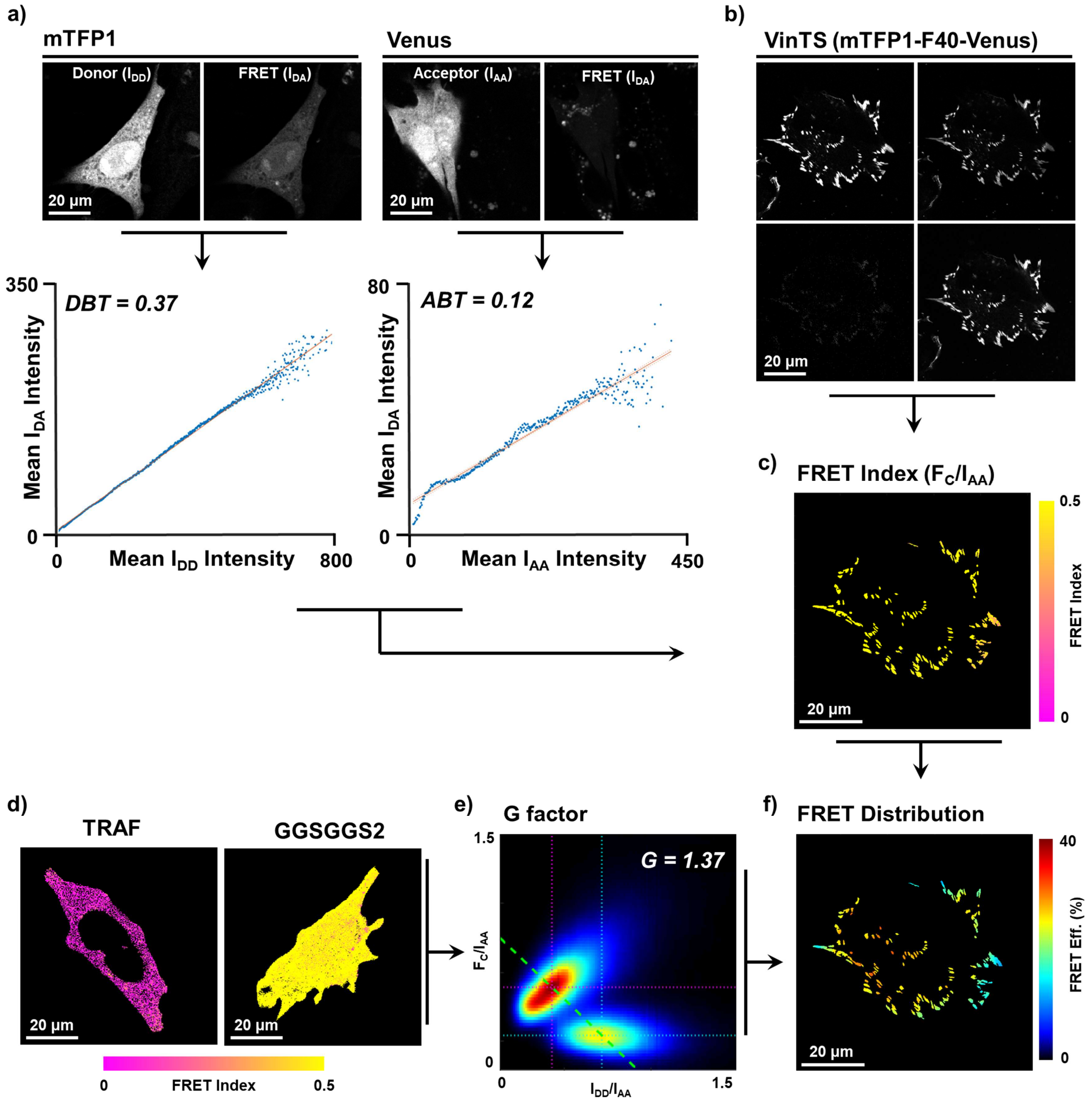


Supplementary Figure 2. Sensitized-emission approach to calculate FRET efficiency of molecular tension probes. (a) Representative cells expressing cytosolic mTFP1 (left) or cytosolic Venus (right). Donor ($I_{DD}$), acceptor ($I_{AA}$), and FRET ($I_{DA}$) channels are shown above. The slopes of the linear regression of mean $I_{DA}$ versus mean $I_{DD}$ or $I_{AA}$ were used to determine donor bleedthrough (DBT = 0.37) and acceptor bleedthrough (ABT = 0.12), respectively. (b) Donor, acceptor, and FRET channel images of a representative cell expressing VinTS with F40 sensor module and mTFP1-Venus as FRET pair. (c) Corresponding FRET index image ($F_C/I_{AA}$) after correction of FRET channel for DBT and ABT and normalization to $I_{AA}$. (d) FRET index images of cells expressing the low-FRET control mTFP1-TRAF-Venus (left) and the high-FRET control mTFP1-$(GGSGGS)_2$-Venus (right). (e) Two-dimensional histogram of $F_C/I_{AA}$ versus $I_{DD}/I_{AA}$ from the low- and high-FRET controls used to determine the G-factor from the slope connecting the distribution peaks (G = 1.37). (f) FRET efficiency map of the VinTS-expressing cell in (b–c) obtained by converting the FRET index using the G-factor. n = 15 cells for mTFP1, Venus, mTFP1-TRAF-Venus, and mTFP1-$(GGSGGS)_2$-Venus samples.

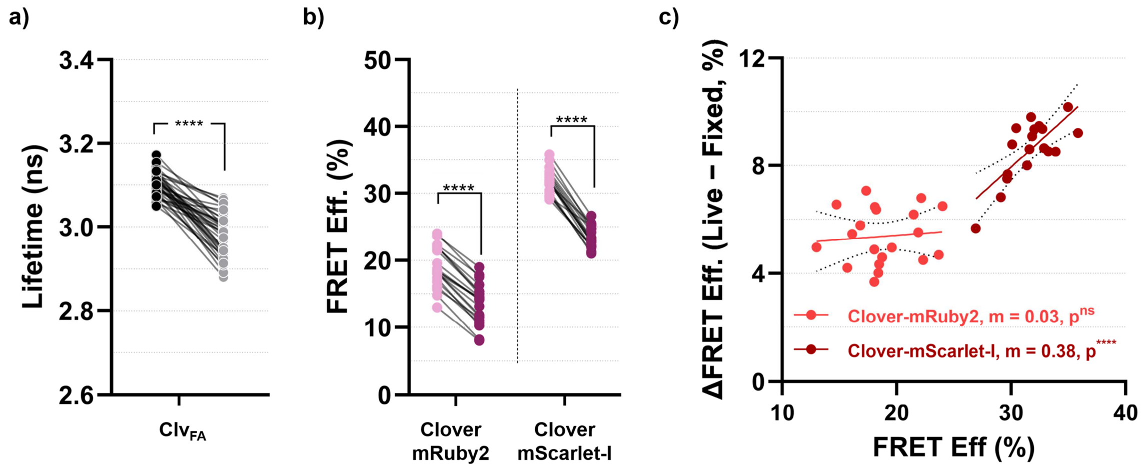


Supplementary Figure 3. Effect of PFA fixation on FRET distribution of F40 mechanical linker with different fluorophore pairs. (a) Fluorescence lifetime of FA-localized Clover ($Clv_{FA}$), generated by positioning Clover between the vinculin head and tail domains, measured in the same cells before and after PFA fixation. Clover lifetime decreased from 3.10 ± 0.02 ns in live cells to 2.98 ± 0.05 ns after fixation (mean paired decrease: 0.12 ns; n = 38 cells). (b) Paired comparison of FRET efficiency in $VinTS^{I997A}$ containing Clover-mRuby2 or Clover-mScarlet-I before and after PFA fixation. FRET efficiency decreased from 19.0 ± 3.0% to 13.5 ± 3.0% for Clover-mRuby2 and from 31.8 ± 1.9% to 23.4 ± 1.6% for Clover-mScarlet-I, corresponding to mean paired decreases of 5.5% and 8.4%, respectively. n = 21 cells for Clover-mRuby2 and 19 cells for Clover-mScarlet-I. (c) Fixation-induced change in FRET efficiency (live − fixed) plotted against the initial live-cell FRET efficiency for the same cells shown in (b). Lines show linear regression fits with 95% confidence intervals; slopes (m) and statistical significance are indicated in the figure. Statistical comparisons in (a) and (b) were performed using paired two-tailed Student's t-tests: ****p < 0.0001.

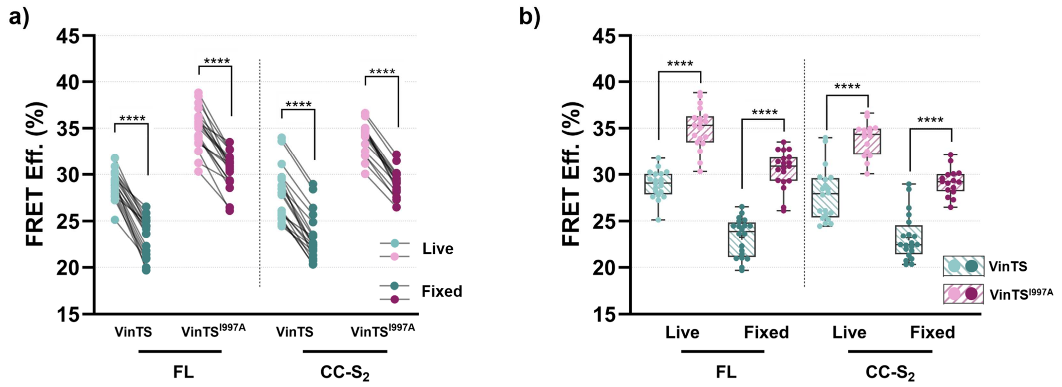


Supplementary Figure 4. Effect of PFA fixation on the FRET response of FL- and CC-S$_2$-based vinculin tension sensors. (a) Paired comparison of FRET efficiency in the same cells before and after PFA fixation for VinTS and VinTS$^{I997A}$ containing FL or CC-S$_2$ as mechanical linker module. For FL, FRET efficiency decreased from 28.9 ± 1.5% to 23.3 ± 2.1% in VinTS and from 35.0 ± 2.3% to 30.6 ± 2.0% in VinTS$^{I997A}$, corresponding to mean paired decreases of 5.7% and 4.5%, respectively. For CC-S$_2$, FRET efficiency decreased from 27.9 ± 2.8% to 23.2 ± 2.5% in VinTS and from 33.8 ± 1.8% to 29.2 ± 1.5% in VinTS$^{I997A}$, corresponding to mean paired decreases of 4.7% for both constructs. (b) Comparison of VinTS and VinTS$^{I997A}$ FRET efficiencies in live and fixed conditions using the same cells shown in (a). The VinTS$^{I997A}$-VinTS difference was 6.1% in live cells and 7.3% in fixed cells for FL, and 6.0% in both live and fixed cells for CC-S$_2$. n = 20 cells for VinTS (FL), 19 cells for VinTS$^{I997A}$ (FL), 20 cells for VinTS (CC-S$_2$), and 16 cells for VinTS$^{I997A}$ (CC-S$_2$). Each data point represents a single cell. Statistical analyses between two groups were performed using paired (a) or unpaired (b) two-tailed parametric Student's t-test: ****$p < 0.0001$.

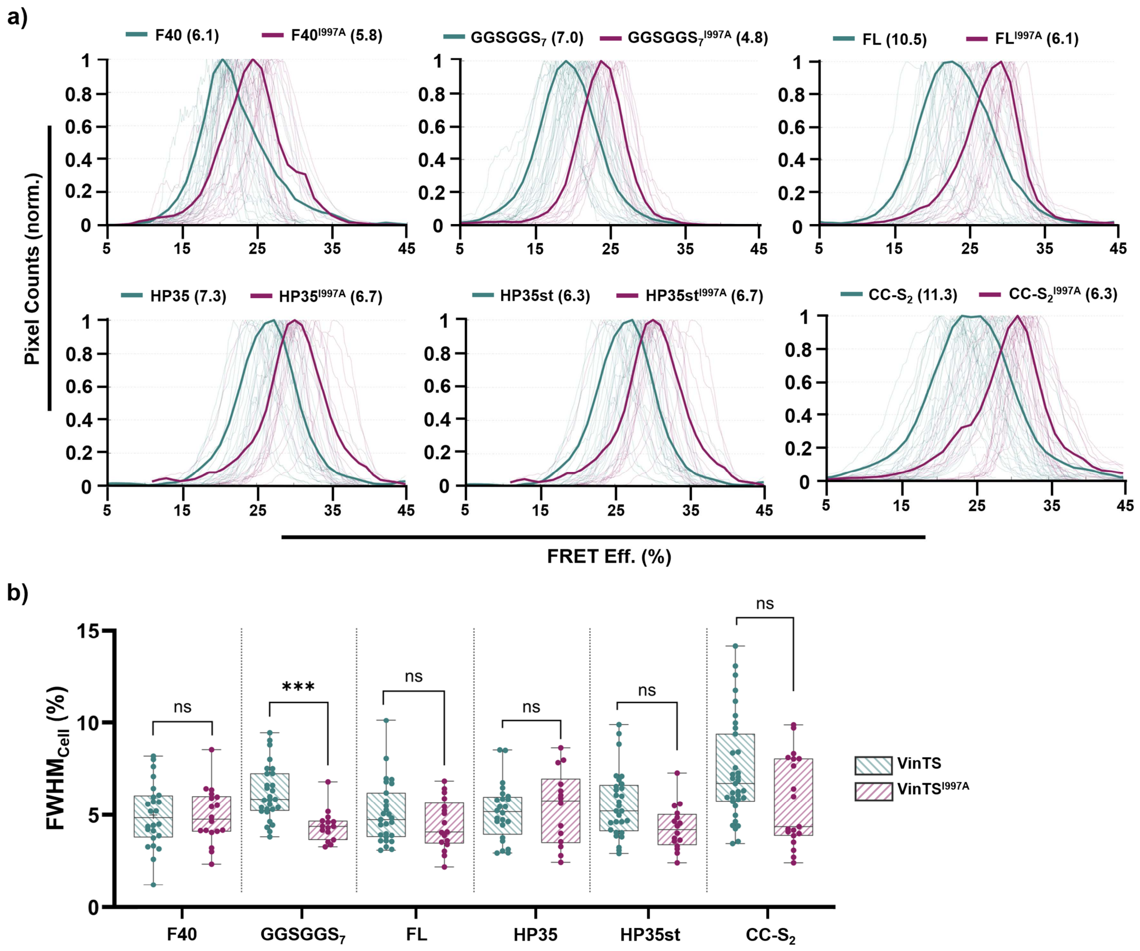


Supplementary Figure 5. Population- and cell-level FRET distribution widths across sensor modules. (a) Normalized FRET efficiency distributions of FA pixels for VinTS (green) and VinTS$^{I997A}$ (purple) containing F40, GGSGGS$_7$, FL, HP35, HP35st, or CC-S$_2$. Thin lines represent individual-cell distributions and thick lines the population-averaged distributions. FWHM values of the population-averaged distributions are indicated in parentheses above each corresponding construct. (b) Per-cell FWHM values for the same VinTS and VinTS$^{I997A}$ cells shown in (a). Each point represents one cell. n = 24, 28, 28, 24, 29, and 35 cells for VinTS and 18, 15, 18, 13, 16, and 20 cells for VinTS$^{I997A}$, respectively. Statistical analyses between two groups were performed using paired unpaired two-tailed parametric Student's t-test: ***p < 0.001.

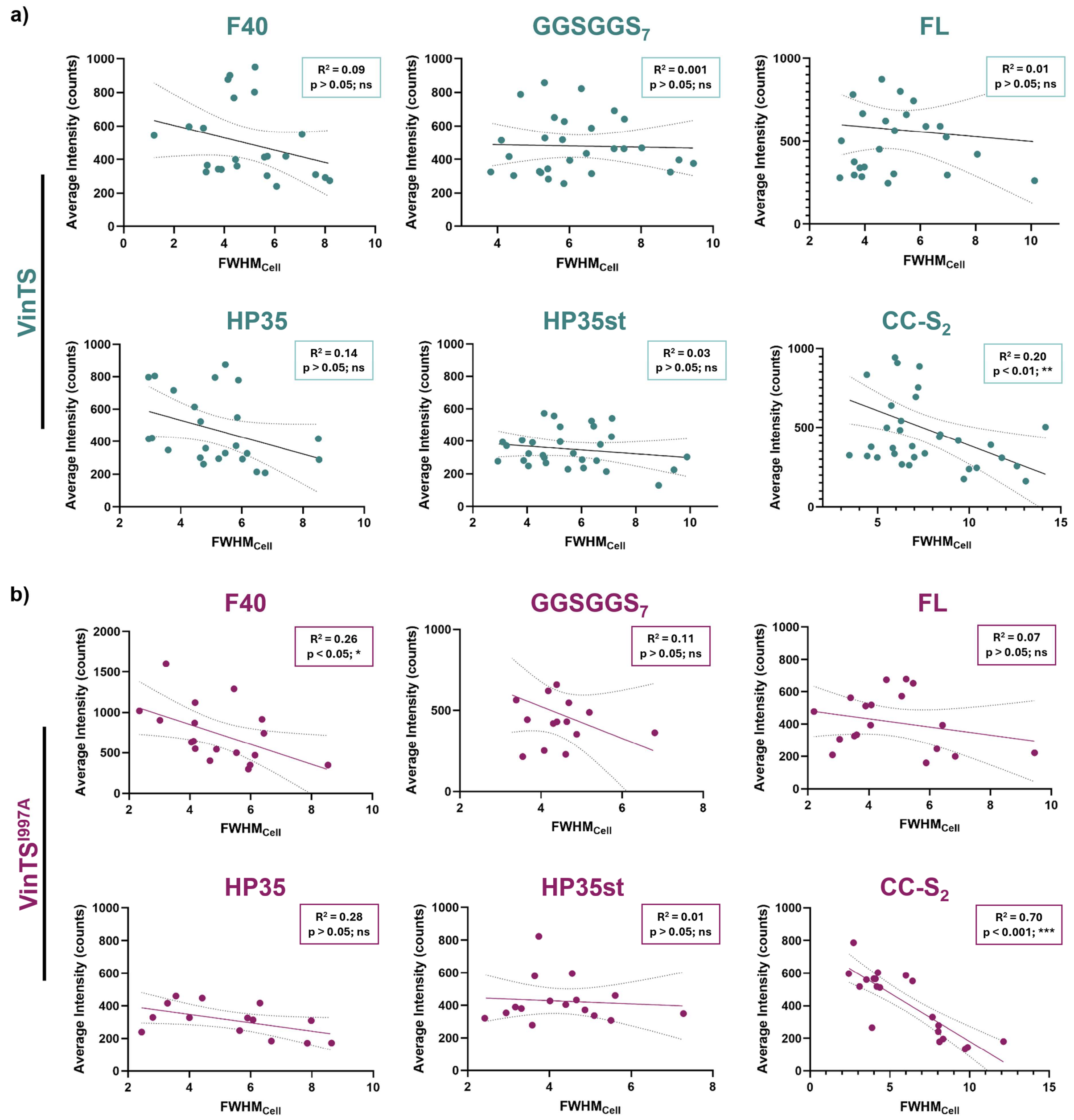


Supplementary Figure 6. Relationship between per-cell FRET distribution width and donor intensity. Average donor-channel intensity within FA masks plotted against per-cell FWHM for (a) VinTS and (b) VinTS$^{I997A}$ across the six sensor modules. The same cells analyzed in Supplementary Fig. 5 were used. Each data point represents a single cell. Solid lines show linear regression fits and dotted lines the corresponding 95% confidence intervals. $R^2$ values and statistical significance of the regressions are indicated in each panel. *$p < 0.05$, **$p < 0.01$, ***$p < 0.001$.

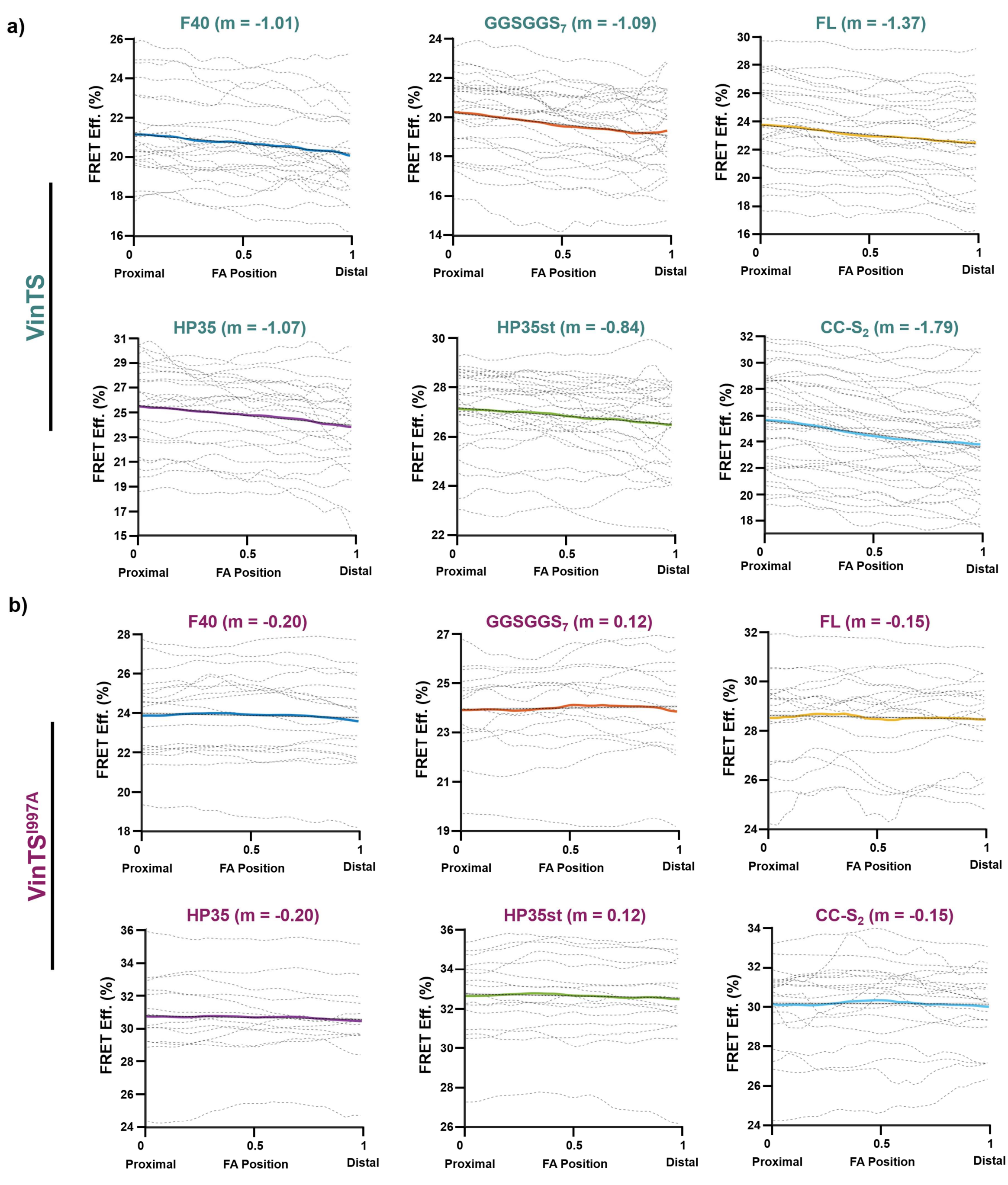


Supplementary Figure 7. Sub-FA FRET distribution of VinTS and VinTS$^{I997A}$ with different sensor modules. (a, b) Spatial FRET-efficiency profiles along the proximal-to-distal end of FAs for cells expressing (a) VinTS or (b) VinTS$^{I997A}$ with distinct sensor modules. Dashed gray lines represent individual cell-averaged profiles, and thick colored lines represent the global average for F40 (blue), GGSGGS$_7$ (orange), FL (yellow), HP35 (purple), HP35st (green), and CC-S$_2$ (cyan). The slope reported above each plot was obtained from linear regression of the corresponding global average line. For VinTS, the analysis included 533 FAs from 24 cells (F40), 796 FAs from 28 cells (GGSGGS$_7$), 640 FAs from 28 cells (FL), 613 FAs from 24 cells (HP35), 783 FAs from 29 cells (HP35st), and 967 FAs from 35 cells (CC-S$_2$). For VinTS$^{I997A}$, the analysis included 573 FAs from 18 cells (F40), 372 FAs from 15 cells (GGSGGS$_7$), 454 FAs from 18 cells (FL), 453 FAs from 13 cells (HP35), 490 FAs from 16 cells (HP35st), and 586 FAs from 20 cells (CC-S$_2$).

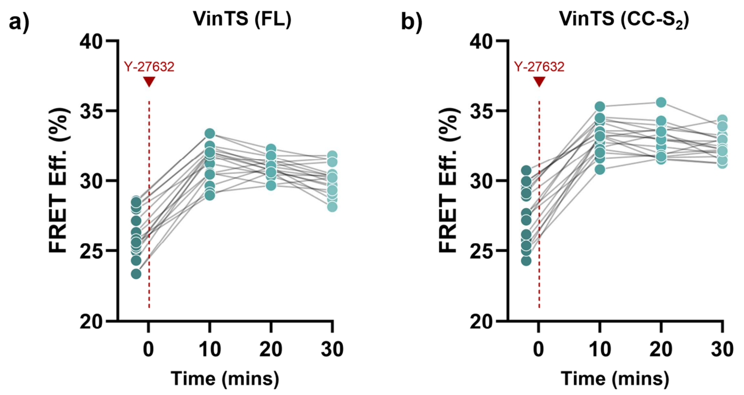


**Supplementary Figure 8. FRET response of FL- and CC-$S_2$-based VinTS following inhibition of ROCK-mediated contractility.** FRET efficiency was monitored in the same live cells expressing VinTS containing (**a**) FL or (**b**) CC-$S_2$ before and after treatment with the ROCK inhibitor Y-27632. Immediately after the initial measurement, Y-27632 was added to a final concentration of 5 µM, and the same cells were imaged again at approximately 10, 20, and 30 min. FRET efficiency increased from 26.1 ± 1.7% to 31.4 ± 1.3% within the first 10 min for FL, corresponding to a mean paired increase of 5.3%, and from 27.6 ± 2.0% to 33.2 ± 1.2% for CC-$S_2$, corresponding to a mean paired increase of 5.6%. Each line connects measurements from the same cell. n = 17 cells for both FL and CC-$S_2$.

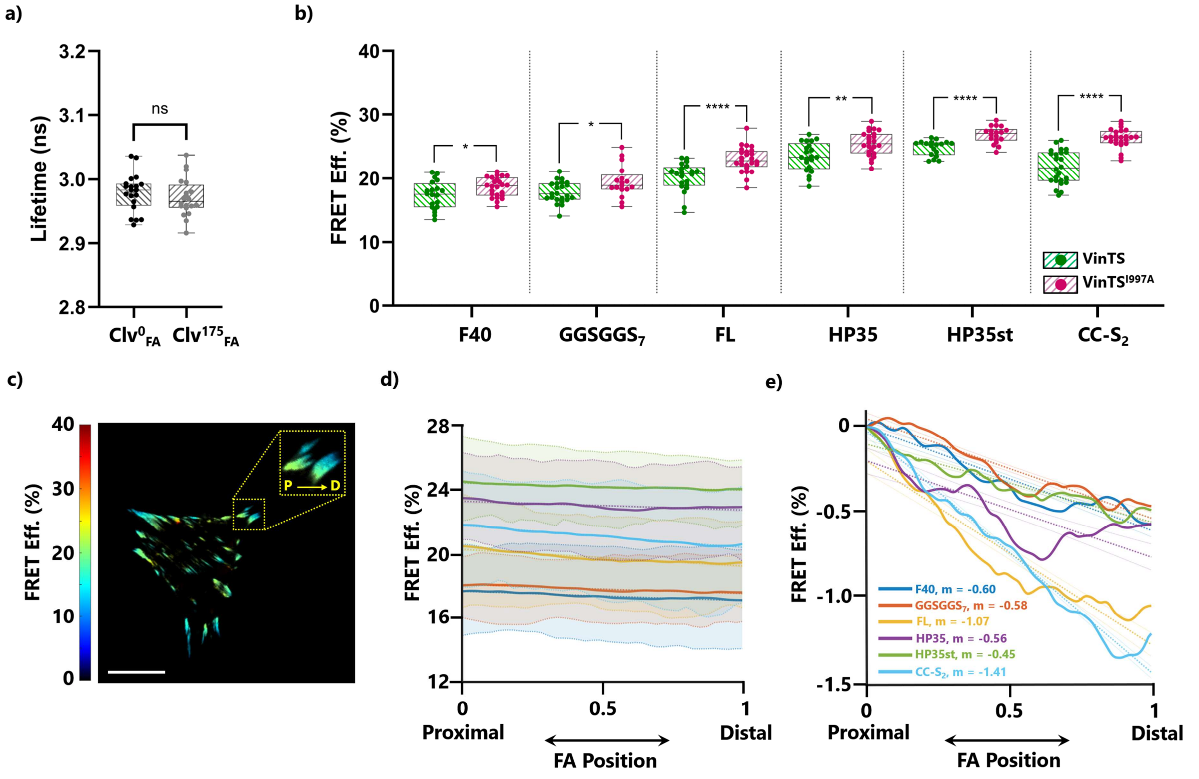


Supplementary Figure 9. Effect of Clover circular permutation on donor lifetime, FRET efficiency, and sub-FA FRET gradients. (a) Fluorescence lifetime of FA-localized $Clv_{FA}^{175}$ compared to FA-localized unpermuted Clover ($Clv_{FA}^{0}$). Each data point represents a single cell. n= 20 cells per condition. (b) Cell-averaged FRET efficiency of VinTS and $VinTS^{I997A}$ constructs containing $Clv^{175}$ and F40, $GGSGGS_7$, FL, HP35, HP35st or CC-$S_2$ as the mechanical sensor domain. Each data point represents a single cell. n = 20, 23, 21, 21, 18 and 24 cells for VinTS; n = 24, 17, 25, 24, 18 and 24 cells for $VinTS^{I997A}$ containing F40, $GGSGGS_7$, FL, HP35, HP35st and CC-$S_2$, respectively. Statistical analyses were performed using unpaired two-tailed Student's t-tests: *$p < 0.05$, **$p < 0.01$, ****$p < 0.0001$. (c) Representative FRET efficiency image showing the FRET gradient across individual FAs in a cell expressing VinTS with $Clv^{175}$ variant and CC-$S_2$ module. Scale bar: 25 µm. (d) Line plots of the spatial FRET efficiency profiles along the proximal-to-distal end of FAs. Shaded regions represent the distribution of cell-averaged data; thick lines represent the global average of VinTS containing F40 (blue), $GGSGGS_7$ (orange), FL (yellow), HP35 (purple), HP35st (green) and CC-$S_2$ (cyan) as the mechanical sensor domain. (e) Baseline-normalized line plots of the global average profiles shown in (f); n = 632 FAs from 20 cells (F40), 573 FAs from 23 cells ($GGSGGS_7$), 577 FAs from 21 cells (FL), 493 FAs from 21 cells (HP35), 469 FAs from 18 cells (HP35st) and 667 FAs from 24 cells (CC-$S_2$) for line plots of (d) and (e). Slopes obtained from linear regression fits of the global average lines.

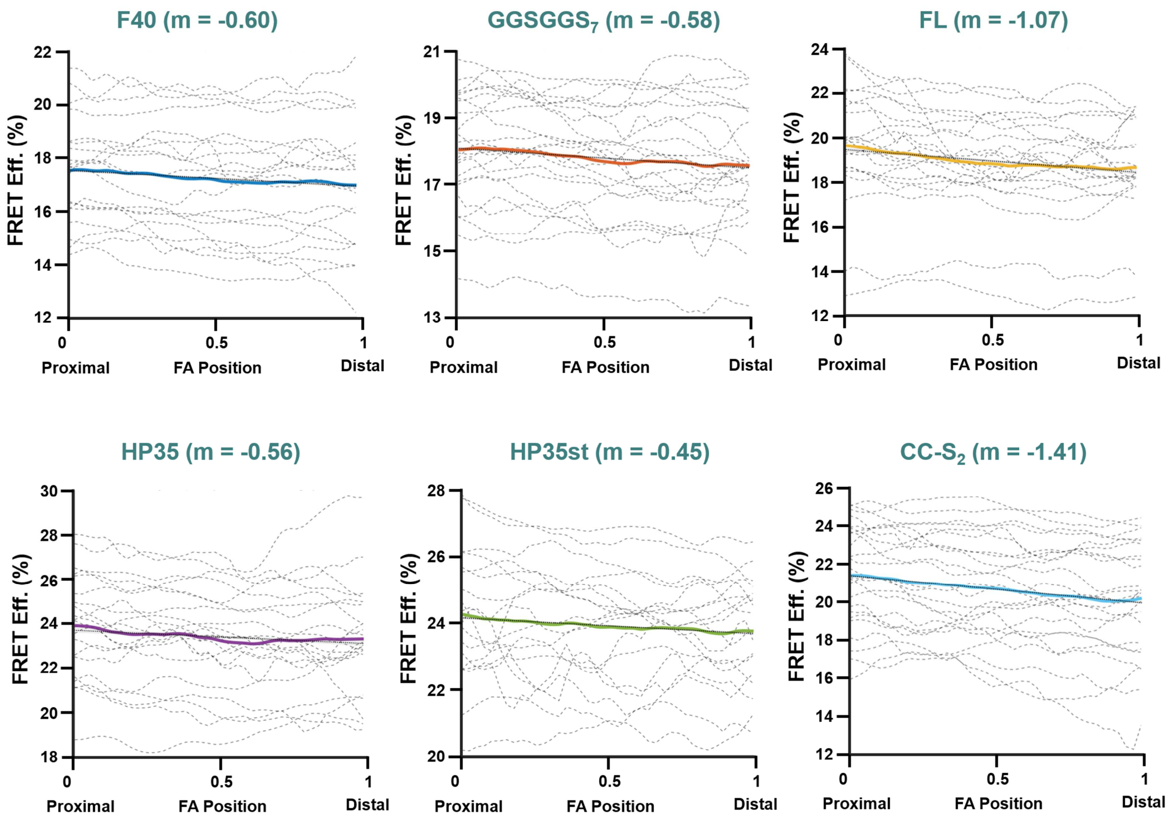


Supplementary Figure 10. sub-FA FRET distribution of VinTS (Clv$^{175}$) for individual sensor modules. (a-f) Spatial FRET efficiency line profiles along the proximal-to-distal end of FAs of cells expressing VinTS with Clv$^{175}$ and distinct sensor modules. Dashed gray lines represent the distribution of cell-averaged data; thick lines represent the global average of VinTS with (a) F40 (blue), (b) GGSGGS$_7$ (orange), (c) FL (yellow), (d) HP35 (purple), (e) HP35st (green) and (f) CC-S$_2$ (cyan) as the mechanical sensor domain. Slope reported above each plot was obtained from the linear regression fits of the corresponding global average line. n = 632 FAs from 20 cells (F40), 573 FAs from 23 cells (GGSGGS$_7$), 577 FAs from 21 cells (FL), 493 FAs from 21 cells (HP35), 469 FAs from 18 cells (HP35st) and 667 FAs from 24 cells (CC-S$_2$).

| Primer ID | Primer Sequence (5'-3') |
|---|---|
| pcDNA-FP-EcoRI | CTGCGGGAATTCTCTAGAGGGCCCGTTTAAACCC |
| pcDNA-RP-XhoI | CTGCGGCTCGAGGGTAAGCTTAAGTTTAAACG |
| VinHD-RP-XhoI | CCATCATCTCGAGCTCATCTTTTTCTTCAGG |
| VinTD-FP-NotI | CCGGTTACAAGGCGGCCGCGGAGTT |
| VinTD-RP-EcoRI-Stop | TCTGCAGAATTCTTACTGATACCATGGG |
| mTPF1-FP-EcoRI | CTGCGGGAATTCATGGTGAGCAAGGGCGAGGAGACCACAATG |
| mTFP1-FP-XhoI | TGTACTCGAGATGGTGAGCAAGGGCGAGGA |
| mTFP1-RP-Stop-Bsp1407I | TCTGCATTACTTGTACAGCTCGTCCATGCCG |
| mTFP1-RP-NotI | TCTGCAGCGGCCGCCTTGTACAGCTCG |
| mTFP1-RP-GGSGGS-Kpn2I | CCGCAGTCCGGATCCACCGGATCCACCCTTGTACAGCTCGTCCATG |
| Venus-RP-GGSGGS-Kpn2I | CCGCAGTCCGGAGGTTCTGGTGGATCCATGGTGAGCAAGGGCGA |
| Venus-RP-EcoRI-Stop-Bsp1407I | CTGCGGGAATTCTTACTTGTACAGCTCGTCCATG |
| Venus-RP-XbaI-Stop-Bsp1407I | CTGCGGTCTAGATTACTTGTACAGCTCGTCCATGCCGAGAGTGATC |
| Clover-FP-XhoI | GATGAGCTCGAGATGATGGTGAGCAAGGGCG |
| Clover-RP-Kpn2I | ATTATATCCGGAGGCGGCGGTCACGAACTCCAGCAG |
| Clover-RP-NotI | TCTGCAGCGGCCGCCTTGTACAGCTCG |
| mRuby2-FP-BamHI | GTGGAGCTGGATCCATGGTGTCTAAGGGCGAAG |
| mRuby2-RP-EcoRI-Stop-Bsp1407I | CTGCGGGAATTCTTACTTGTACAGCTCGTCCATG |
| mScarlet-I-FP-BamHI | GTGGAGCTGGATCCATGGTGTCTAAGGGCGAAG |
| mScarlet-I-RP-EcoRI-Stop-Bsp1407I | CTGCGGGAATTCTTACTTGTACAGCTCGTCCATG |
| TRAF-FP-Kpn2I | CTGCGGTCCGGAGAGAGCCTGGAGAAGAAGACGGCCACTTT |
| TRAF-RP-BamHI | CTGCGGGGATCCGAGCCCTGTCAGGTCCACAATGGCCTTGAT |
| Clover-[176-236]-FP-XhoI | CTTACCCTCGAGATGAGCGTGCAGCTCGCCGAC |
| Clover-[176-236]-RP-GGTGGS | AGAACCCCCGGTACCTCCCTTGTACAGCTCGTCCAT |
| Clover-[1-175]-FP-GGTGGS | GGAGGTACCGGGGGTTCTATGGTGAGCAAGGGCGAGGAGCTG |
| Clover-[1-175]-RP-Kpn2I | TCTAGATCCGGAGCCGTCCTCAACGTTGTGGCGGATC |
| Clover-[1-175]-RP-NotI | GAACTCCGCGGCCGCGCCGTCCTCAACGTTGTGGC |
| Clover-[1-175]-RP-EcoRI-Stop | TCTAGAGAATTCTTAGCCGTCCTCAACGTTGTGGCGGATC |

**Supplementary Table 1. List of primers used to generate the plasmid constructs in this study.** Primer IDs indicate the orientation of the primer (FP, forward primer; RP, reverse primer), the specific restriction cut(s) and the stop codon, if present. All primer sequences are given in 5'-3' direction. Restriction sites are indicated in bold green. Stop codons are underlined.

| gBlock ID | gBlock Sequence (5'-3') |
|---|---|
| F40 (codon optimized) | AACGAGAAGCGCGATCACATGGTCCTGCTGGAGTTCGTGACCGCTGCTTCCGGAGGTCCGGGAGGTGCTGGCCCCGGTGGTGCTGGTCCTGGTGGTGCAGGACCTGGTGGAGCAGGGCCTGGAGGGGCTGGACCCGGGGGTGCGGGTCCTGGAGGAGCCGGTCCAGGTGGAGCTGGATCCAAGGGCGAAGAGCTGATCAAGGAAAATATGCGTATGAAGGTGGTCATGGAA |
| GGSGGS$_7$ (codon optimized) | AACGAGAAGCGCGATCACATGGTCCTGCTGGAGTTCGTGACCGCTGCTTCCGGAGGATCAGGAGGTTCTGGGGGTAGCGGTGGCAGCGGAGGTAGTGGCGGCTCTGGCGGTAGTGGGGGAAGTGGAGGTTCCGGCGGTTCTGGAGGCTCAGGCGGCTCCGGTGGGTCTGGGGGATCCAAGGGCGAAGAGCTGATCAAGGAAAATATGCGTATGAAGGTGGTCATGGAA |
| FL | AACGAGAAGCGCGATCACATGGTCCTGCTGGAGTTCGTGACCGCTGCTTCCGGAATGGGCGAGTTTGACATCCGGTTTCGGACTGATGACGACGAACAGTTCGAGAAAGTGCTGAAGGAGATGAATCGTCGAGCCAGAAAGGATGCTGGAACTGTGACCTACACAAGGGATGGGAATGACTTCGAGATTCGCATTACCGGCATAAGCGAGCAAAACCGCAAAGAACTGGCCAAAGAGGTTGAAAGGCTTGCAAAGGAACAGAACATCACAGTCACGTATACCGAGAGAGGTTCCCTCGAAGGATCCAAGGGCGAAGAGCTGATCAAGGAAAATATGCGTATGAAGGTGGTCATGGAA |
| HP35 | AACGAGAAGCGCGATCACATGGTCCTGCTGGAGTTCGTGACCGCTGCTTCCGGACTGAGCGATGAGGACTTCAAAGCTGTGTTTGGCATGACCAGGTCCGCATTTGCCAATCTTCCTCTGTGGAAACAACAGAACCTGAAGAAGGAAAAGGGACTCTTCGGATCCAAGGGCGAAGAGCTGATCAAGGAAAATATGCGTATGAAGGTGGTCATGGAA |
| HP35st | AACGAGAAGCGCGATCACATGGTCCTGCTGGAGTTCGTGACCGCTGCTTCCGGACTGAGCGATGAGGACTTCAAAGCTGTGTTTGGCATGACCAGGTCCGCATTTGCCAATCTTCCTCTGTGGAAACAACAGGCCCTGATGAAGGAAAAGGGACTCTTCGGATCCAAGGGCGAAGAGCTGATCAAGGAAAATATGCGTATGAAGGTGGTCATGGAA |
| CC-S$_2$ | AACGAGAAGCGCGATCACATGGTCCTGCTGGAGTTCGTGACCGCTGCTTCCGGAGCCCAATTAGAAAAGGAGCTGCAGGCACTTGAGAAAGAGAATGCTCAGTTAGAGTGGGAACTTCAAGCATCGGAGAAGGAATCGGCACAGGGCGGTAGTGGCGGCAGCCAAGCGTCGAAAAAGAAATCGGCGCAGCTGAAATGGAAGTTGCAGGCCAATAAAAAGAAGCTGGCTCAACTGAAGAAAAAGTTGCAAGCCGGATCCAAGGGCGAAGAGCTGATCAAGGAAAATATGCGTATGAAGGTGGTCATGGAA |

Supplementary Table 2. List of gBlock sequences of the sensor modules used in this study. gBlock ID indicates the mechanical sensor module, with codon optimized modules indicated in parentheses. All sequences are given in 5'-3' direction. Kpn2I cut site is indicated in green; BamHI cut site is indicated in red.
